# Single cell profiling of immature human postnatal thymocytes resolves the complexity of intra-thymic lineage differentiation and thymus seeding precursors

**DOI:** 10.1101/2020.04.07.007237

**Authors:** Marieke Lavaert, Kai Ling Liang, Niels Vandamme, Jong-Eun Park, Juliette Roels, Monica S. Kowalczyk, Bo Li, Orr Ashenberg, Marcin Tabaka, Danielle Dionne, Timothy L. Tickle, Michal Slyper, Orit Rozenblatt-Rosen, Bart Vandekerckhove, Georges Leclercq, Aviv Regev, Pieter Van Vlierberghe, Martin Guilliams, Sarah A. Teichmann, Yvan Saeys, Tom Taghon

## Abstract

During postnatal life, thymopoiesis depends on the continuous colonization of the thymus by bone marrow derived hematopoietic progenitors that migrate through the bloodstream. In human, the nature of these thymus immigrants has remained unclear. Here, we employ single-cell RNA sequencing on approximately 70.000 CD34^+^ thymocytes to unravel the heterogeneity of the human immature postnatal thymocytes. Integration of bone marrow and peripheral blood precursors datasets identifies several putative thymus seeding precursors that display heterogeneity for currently used surface markers as revealed by CITEseq. Besides T cell precursors, we discover branches of intrathymic developing dendritic cells with predominantly plasmacytoid DCs. Trough trajectory inference, we delineate the transcriptional dynamics underlying early human T-lineage development from which we predict transcription factor modules that drive stage-specific steps of human T cell development. Thus, our work resolves the heterogeneity of thymus seeding precursors in human and reveals the molecular mechanisms that drive their *in vivo* cell fate.

## Introduction

T cells are central and essential mediators of the adaptive immune system. T cell deficiencies result in life-threatening conditions, such as perturbed and delayed T cell reconstitution following chemo- and radiotherapy. In addition to their protective role against pathogens, T cells are increasingly exploited for immunotherapeutic purposes, particularly through the generation of CAR T-cells. To facilitate and optimize their generation and to enhance T cell recovery in patients, insights into the molecular processes that drive early T cell development after birth are of critical importance.

T lymphocytes require the specific microenvironment of the thymus for their development. Maintenance of physiological thymopoiesis requires the continuous colonization of a small amount of bone marrow (BM) derived T cell precursors whose identity is still unclear in human. The thymic microenvironment gradually converts these immigrating and still multipotent progenitor cells to become T-lineage committed, and also induces extensive proliferation to increase the pool of T cell precursors. Notch1 activation in these T cell progenitors is one of the main drivers of these early events and is initiated through interaction with Notch1 ligands such as DLL4 expressed by the thymic epithelial cells. Experimental mouse models have been extensively used to study the early stages of T cell development, but extrapolation of these insights to human biology has proven to be difficult due to species specific differences. This is clearly exemplified by the differences in cell surface markers that are used to phenotype the most immature T cell precursors and this has obscured a clear comparison of these initial stages of T cell development across species. Also functional perturbation experiments have suggested that the molecular mechanisms driving early T cell development in human might differ compared to in mouse (Ha et al., 2017; Taghon et al., 2009; Van de Walle et al., 2009, 2016).

Immature human thymocytes are characterized by the human stem/progenitor cell marker CD34. While CD1a upregulation within these CD34^+^ thymocytes has been used to mark T-lineage commitment, recent work has proposed that the commitment process is already initiated within the most immature CD34^+^CD1a^-^ thymocytes and marked by the loss of CD44 expression within that subset (Canté-Barrett et al., 2017). In each case, the uncommitted fraction of human thymocytes has an unknown heterogeneity in which a small fraction should represent the BM-derived immigrating T cell precursors. Multiple thymus seeding populations have been reported based on differential CD7 expression and these have led to conflicting reports on whether CD34^+^CD7^-^ (Hao et al., 2008; Six et al., 2007) or CD34^+^CD7^int^ (Haddad et al., 2006) precursors reflect the thymus-seeding progenitors. Also CD34^+^CD38^-^ thymocytes have been proposed to comprise the most immature precursor subset (Dik et al., 2005; Weerkamp et al., 2006). Furthermore, it is unclear in human which other hematopoietic lineages develop intrathymically. The developmental potential of immature human thymocyte subsets has mainly been studied *in vitro* but does not necessarily correlate with actual *in vivo* differentiation. The lack of genetic lineage tracing experiments in human have prevented clear insights into this.

The recent emergence of single cell technologies has provided new opportunities to address the issues mentioned above. To obtain a high-resolution perspective on the heterogeneity and *in vivo* lineage differentiation of human precursors cells in the postnatal thymus, we performed scRNAseq and CITEseq on 70.000 CD34^+^ thymocytes. Through integration with BM and peripheral blood data, we reveal distinct precursor subsets that can seed the thymus which show heterogeneity for the currently used cell surface markers and which show clear T- and DC-lineage differentiation. By also unraveling the precise and kinetic transcriptional changes that drive early human T cell development, this study clarifies the initiating hematopoietic differentiation events that occur in the human postnatal thymus.

## Results

### Single cell transcriptome profiling of immature human thymocytes

To dissect the cellular heterogeneity and transcriptional landscape of immature human thymocytes, we performed scRNAseq on CD34 enriched postnatal thymocytes that were further purified using three different sort layouts (**Figure 1A**). To avoid any potential bias, we sorted the total uncommitted lin^-^CD34^+^CD1a^-^thymocyte fraction from 3 independent donors (Sort 1, **Figures 1A and S1A**). To enrich for the more primitive lin^-^CD34^+^CD7^-^ and lin^-^CD34^+^CD7^int^ subsets that both have been proposed to be the thymus seeding precursors (Haddad et al., 2006; Hao et al., 2008; Six et al., 2007), and to include committed T cell precursors to facilitate tracing of differentiating T-lineage cells, the lin^-^CD34^+^CD7^-/int^, lin^-^CD34^+^CD1a^-^ and lin^-^CD34^+^CD1a^+^ populations from 2 donors were sorted and pooled (Sort 2, **Figures 1A and S1B**). CITEseq was performed on these samples to capture the cell surface expression data for CD7, CD44, CD10 and CD123. Finally, to further enrich for the most immature thymocyte subsets and to include lineage positive cells to improve developmental directionality of non T-lineage subsets, we exploited CD44 as a maker for uncommitted thymocytes (Canté-Barrett et al., 2017) and more mature lineage positive cells (Márquez et al., 1995) to sort the lin^+^, lin^-^CD44^+^CD7^-/+^ and lin^-^CD44^int^CD7^+^ populations from 2 donors (Sort 3, **Figures 1A and S1C**). Overall, this resulted in 7 scRNAseq libraries that span 5 donors with approximately 70.000 cells that represent the entire fraction of human CD34^+^ thymocytes.

**Figure 1:**
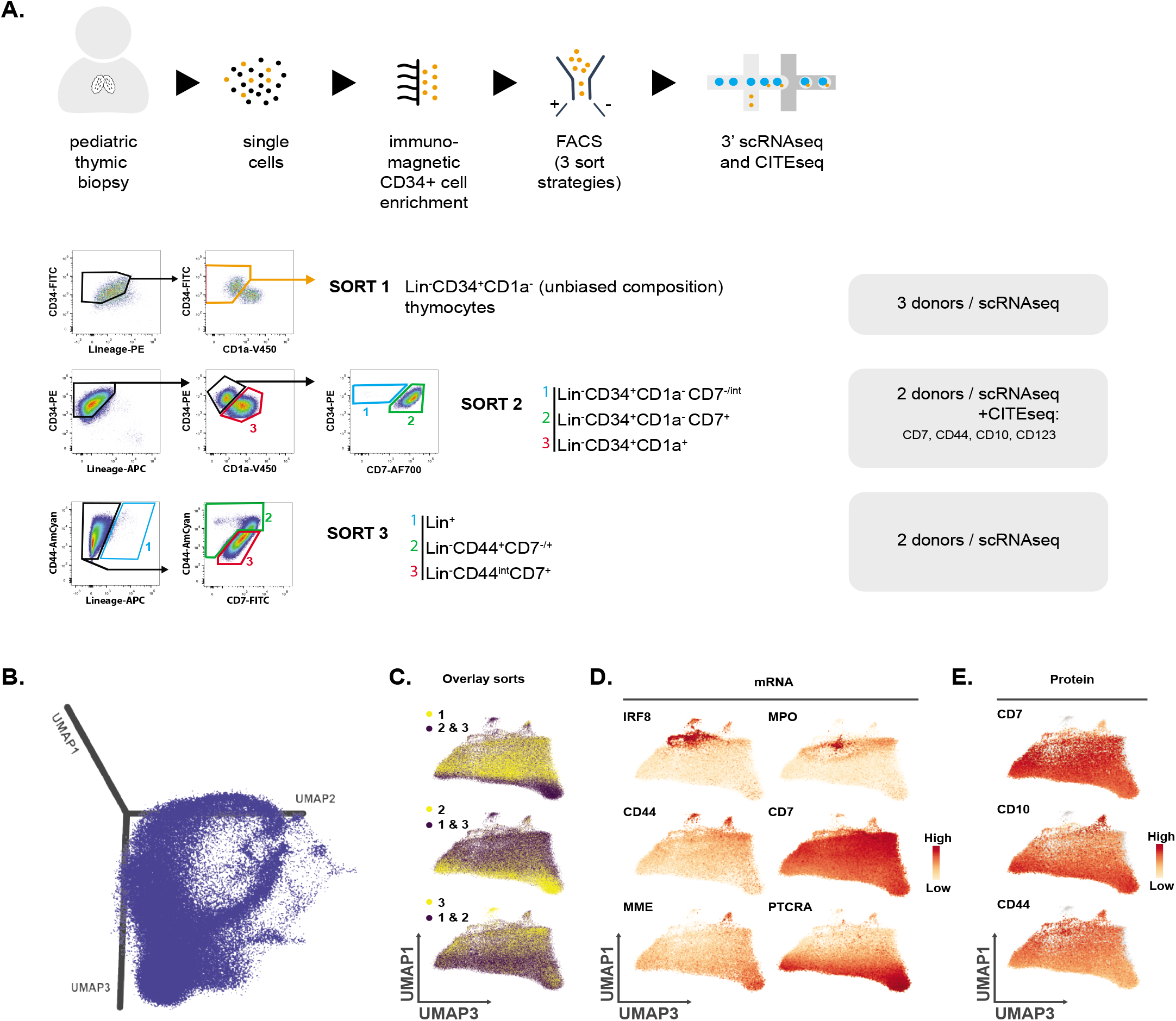
Single cell transcriptome profiling of immature human thymocytes. **A.** Schematic overview of the sample processing pipeline and the 3 different sorting strategies (sort 1-3) used to enrich for rare cell populations. Dot plots show representative sort layouts and 5 different donors were profiled. **B.** 3D UMAP visualization of the *ex vivo* sorted immature human thymocyte dataset. **C.** UMAP visualization of dimensions 1 & 3 shows overlay of each individual sort compared to the other sorting strategies. **D.** UMAP representing the expression of genes associated with either myeloid or early T cell development visualized on dimensions 1 & 3. The size of dots corresponds to gene expression levels. **E.** UMAP representing protein surface expression of CD7, CD10 and CD44 through CITEseq (sort 2), visualized on dimensions 1 & 3. Missing values (sorts 1 & 3) are visualized in grey. The size of dots corresponds to gene expression levels.

On average, we detected between 1500^-^3000 genes per cell, indicating high quality libraries. Low quality cells with spurious mitochondrial mRNA or low sequencing depth, as well as contaminating mature T cells were removed after annotation as described below (**Fig S1D**). To visualize the transcriptional changes, a 3-dimensional UMAP (McInnes et al., 2018) was generated (**Figure 1B**). T cell differentiation seems to largely contribute to variation in the first dimension (UMAP1) since along this axis, the anticipated biological differences between the various sorts are reflected (**Figure 1C**) and a continuous gene expression pattern that is consistent with known changes during mouse and human T cell development can be observed (CD44, CD7, MME and PTCRA, **Figure 1D**) (Canté-Barrett et al., 2017; Hao et al., 2008; Taghon et al., 2009; Van de Walle et al., 2009). Cells expressing high levels of *IRF8* and *MPO* were clearly distinct from the main thymocyte population that expressed CD7 and therefore most likely represent non T-lineage cells (Plum et al., 1999). We also identified cells with intermediate expression of *IRF8* and *MPO*, corresponding to immature T cell precursors as identified by the strong and Notch-dependent expression of *CD7* (Van de Walle et al., 2009) and *CD44* (*García-Peydró et al., 2018*). The expression of these genes decreased upon further T-lineage differentiation towards *PTCRA* expressing thymocytes and coincided with upregulation of *MME* which encodes for CD10 (**Figure 1D**). CITEseq analysis revealed a consistency between RNA and protein expression for CD7, CD44 and CD10, thereby validating the robustness of our transcriptome profiling (**Figure 1E**).

To facilitate the annotation of our CD34^+^ thymocytes, we integrated our data with the human cell atlas (HCA) bone marrow (BM) dataset that provides a development framework to identify all the various cell types within our dataset. Cell clusters were annotated based on the expression of selected marker genes from literature (**Figure 2A**). This was further supported by hierarchical clustering which confirmed that similar cell types cluster more closely together (**Figure 2B**). Besides non T-lineage cells, identified by the expression of genes including *IRF8, MPO, SPI1, CD79B* and *CD19*, we identified a cluster of putative thymus seeding precursors (TSP) based on the expression of the anticipated chemokine receptors *CCR7* and *CCR9* (Krueger et al., 2010; Zlotoff et al., 2010) and the hematopoietic progenitor genes (*BCL11A* and *HOXA9*) and *CD44* (**Figure 2C**) (Yui and Rothenberg, 2014). Consistent with the multipotent potential of the most immature thymocytes, these TSPs clustered together with hematopoietic stem cells (HSCs) and hematopoietic progenitor cells (HPCs) in the BM (**Figure 2B**); and expressed *MME* (**Figure 2C, 1D**) that encodes CD10 (**Figure 1E**), consistent with literature (Six et al., 2007). Early T cell progenitors (ETP) displayed characteristics of strong Notch activation (*CD7*) and clustered clearly distinct from HSCs and HPCs (**Figure 2B**) but still expressed hematopoietic progenitor genes such as *CD34* and *LYL1*. These genes were gradually downregulated as T-cell lineage specification and commitment progressed (**Figure 2C**). This differentiation pattern corresponded with an increase in the expression of critical T-lineage transcription factors (*TCF7, GATA3, BCL11B*) and the immature thymocyte marker *CD1A*, resulting in thymocytes that initiate TCR rearrangements (*DNTT, RAG2, PTCRA*) (**Figure 2C**). The validity of our inferred developmental stages was further supported both by their spatial localization along UMAP1 (**Figure 2D**) and by the developmental trajectory inferred by RNA velocyto (**Figure S2A**). In addition to differentiation, proliferation is also a major biological event during early T cell development. Consistently, UMAP 2 and 3 reflected the variation in expression of specific genes (**Figure 2E**) that corresponded with annotated cell cycle stages (**Figures 2F, S2B-D**) and this was further confirmed by RNA velocyto (**Figure S2E**) (Macosko et al., 2015).

**Figure 2:**
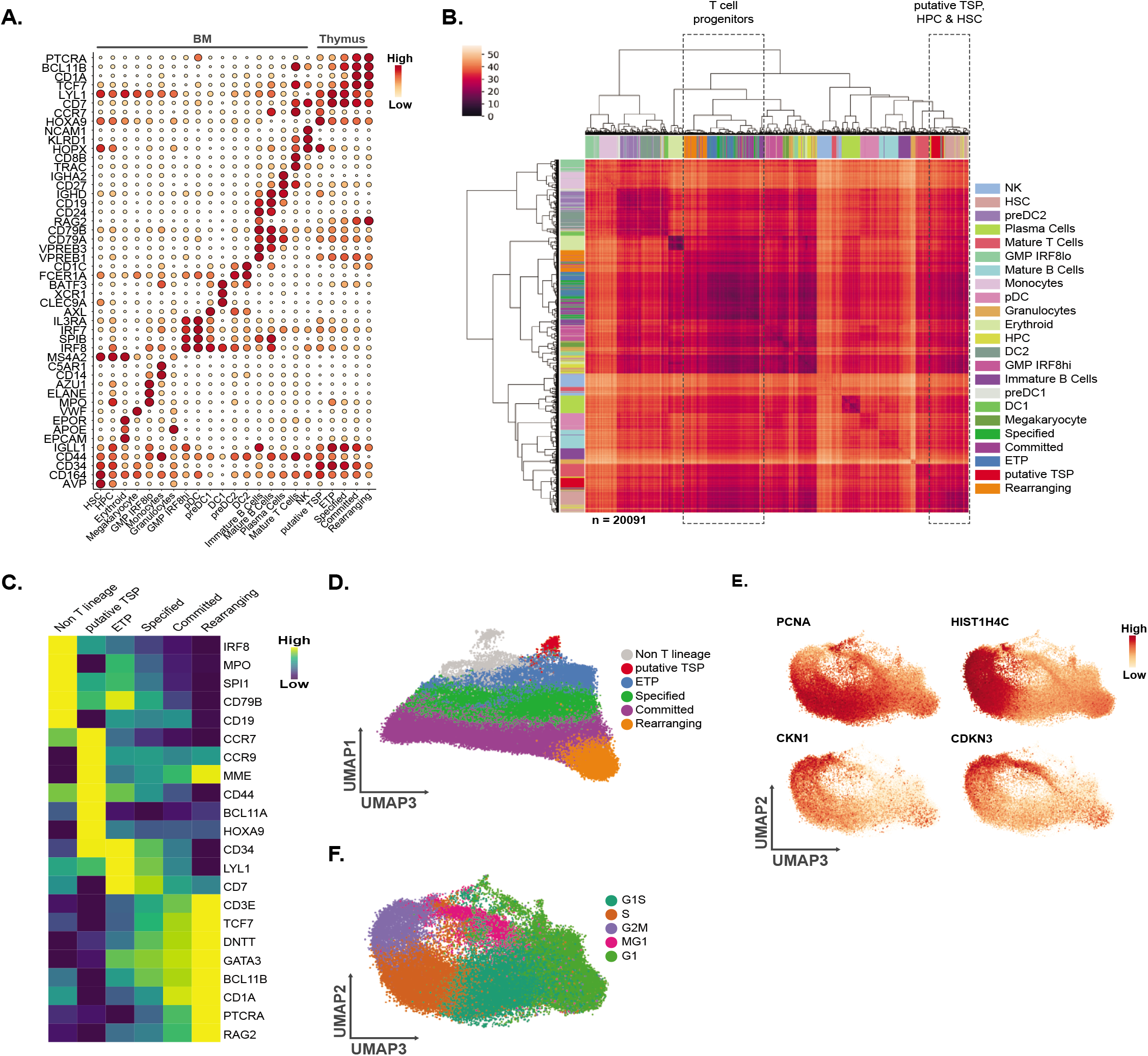
Cellular composition of human CD34+ postnatal thymocytes. **A.** Dot Plot visualizing pseudo bulk log2 counts, averaged and scaled gene-wise, with both color and size corresponding to expression levels of subset specific genes for both the HCA BM and *ex vivo* thymocyte datasets that were both generated using the 10x genomics V2 kit. Detailed information on cell type annotation can be found in the material and methods section. **B.** Distance matrix containing randomly subsampled cells (n = 20091) from both the HCA BM and *ex vivo* thymocyte dataset, which were hierarchically clustered (ward). **C.** Heatmap showing pseudo bulk log2 counts, averaged and scaled gene-wise, of selected lineage- and/or stage-specific genes within annotated CD34^+^ thymocyte subsets. **D.** UMAP visualization of the annotated populations displayed on the dimensions 1 & 3. **E.** UMAP representing the expression of cell cycle associated genes on dimensions 2 & 3. The size of dots corresponds to gene expression levels. **F.** UMAP visualization of the annotated cell cycle stages displayed on dimensions 2 & 3.

### Validation and transcriptional characterization of human thymus seeding precursors

Postnatal thymopoiesis is dependent on the continuous colonization of the thymus by bone marrow derived progenitor cells (**Figure 3A**). Therefore, we anticipated that transcriptionally similar cells would be present in both our immature thymocyte dataset and the published HCA BM dataset, and that these might represent the human thymus seeding precursors whose identity has remained unclear thus far. Using the integrated datasets as described above, we could clearly distinguish the various blood cell lineages and precursor subsets (**Figure S3A**). From this, we retained the CD34^+^ BM cells and the immature CD34^+^ G1 thymocytes (**Figure S3B**) that do not display a thymus specific gene expression pattern (**Figure S3C**) to obtain a high resolution snapshot of thymus and BM resident CD34^+^ precursor cells (**Figure 3B**) that show clear overlap (**Figure 3C**). In addition to other precursor subsets that will be discussed later in the manuscript, clustering algorithms allowed us to identify the BM resident cells that consistently grouped together with the thymic putative TSPs (Cluster 8 and 16, **Figure S3D**), resulting in the identification of 44 BM-resident TSPs (**Figure S3E**). Interestingly, these populations revealed a similar topological location in the UMAP prior to and after migration into the thymus (**Figure 3B** and **Figure S3E**). Consistent with our initial annotation (**Figure 2C**), these putative intra-thymic TSPs were CD7 negative and *MME/*CD10 positive at both the protein and mRNA level (**Figure 3D-E**), and expressed both *CD34* and *CD44*, but lacked abundant expression of genes associated with myeloid (*IRF8, MPO*), erythroid (*GATA1*) or B-lineage (*IGLL1*) cells compared to other, more lineage restricted precursor subsets (**Figure 3F and Figure S3F**). Differential gene expression analysis across the populations, retaining only genes that showed consistent variation in both the BM dataset and the integrated BM-thymus datasets to remove thymus-specific genes, revealed a unique transcriptional signature for the putative TSPs (**Fig 3G**), including expression of genes such as *LTB*, *EVL, HOPX, KIAA0087, CCR7, HOXA9, HOXA10, ARL6IP5* and *KIAA0125* in both the BM and thymus resident TSPs (**Figure S3G-H**).

**Figure 3:**
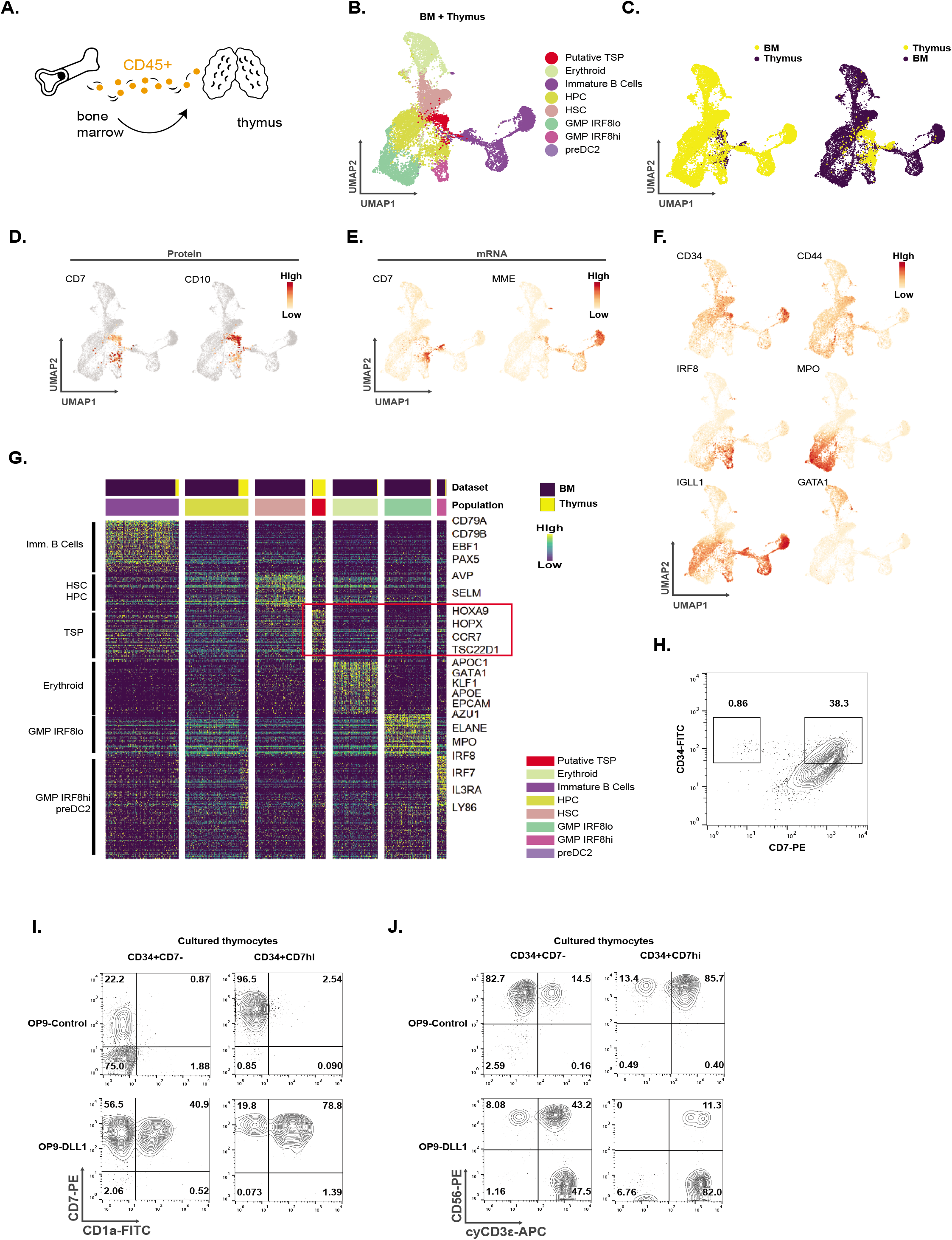
Validation and characterization of the human thymus seeding progenitor (TSP). **A.** Schematic overview of the conceptual migration of a putative TSP from the BM into the thymus to initiate thymopoiesis. **B.** UMAP visualization of the annotated CD34^+^ BM cells and overlapping CD34^+^ G1 thymocyte populations following integration as described in **Figure S3A-C**. **C.** UMAP visualization showing the overlay of the integrated and retained CD34^+^ BM cells and CD34^+^ G1 thymocytes. **D.** UMAP representing protein surface expression of CD7 and CD10 through CITEseq (sort 2) on the CD34^+^ G1 thymocytes that overlap with BM CD34^+^ cells. Missing values (sorts 1 & 3) are visualized in grey. The size of dots corresponds to gene expression levels. **E-F.** UMAP representing the expression of selected genes in the integrated and retained CD34+ BM cells and CD34+ G1 thymocytes. The size of dots corresponds to gene expression levels. **G.** Heatmap displaying differentially expressed genes in the various annotated subsets that are present in the integrated and retained CD34^+^ BM cells and overlapping CD34+ G1 thymocytes. **H.** Contour plot showing representative CD34 versus CD7 flow cytometric analysis in *ex vivo* isolated lin^-^CD34^+^ thymocytes. Numbers in the contour plots display the frequency of the CD7^-^ and CD7^hi^ populations. **I-J.** Flow cytometric analysis of *ex vivo* isolated lin^-^CD34^+^CD7^-^ and lin^-^CD34^+^CD7^hi^ thymocytes following coculture on either OP9-control (top row) or OP9-DLL1 (bottom row) stromal cells. Density plots show CD7 versus CD1a staining to reveal T-lineage potential (**I**) or CD56 versus intracellular CD3ε staining to reveal Notch priming (**J**) following 2 weeks of co-culture. Numbers in the density plots display the frequencies of the corresponding populations. Data are representative for at least 3 independent experiments.

Although the CD34^+^CD7^-^CD10^+^ phenotype of our putative TSP is based on both mRNA (**Figure 3D**) and protein data (**Figure 3E**), and also consistent with earlier studies (Hao et al., 2008; Six et al., 2007), we further confirmed the presence of a rare CD34^+^CD7^-^ population (**Figure 3H**) within the CD34^+^ thymocyte fraction that displays clear potential to differentiate into CD7^+^CD1a^+^ committed T-lineage cells, but with slower kinetics compared to Notch-dependent CD34^+^CD7^hi^ thymic progenitors in agreement with a more primitive developmental state (**Figure 3I**). Using expression of intracellular CD3ε as a hallmark of Notch activation (De Smedt et al., 2007), these CD34^+^CD7^-^ putative TSPs displayed few signs of Notch signaling and lacked expression of *CD3E* (**Figure 2C**), and mainly generated CD56^+^CD3ε^-^ NK cells on OP9-control stromal cells, in contrast to CD34^+^CD7^hi^ thymocytes that predominantly differentiated into CD3ε expressing NK cells in the absence of further Notch activation (**Figure 3J**). Both precursor subsets generated CD3ε^+^ NK- and T-lineage cells on OP9-DLL1 stromal cells, illustrative of their capacity to respond to Notch activation (**Figure 3I-J**). These findings thus suggest that the CD34^+^CD7^-^ thymocytes are Notch^-^naive T cell progenitors that represent the most immature cells that have just entered the human thymus.

### Transcriptional dynamics of early human T cell development

In human, the precise transcriptional events that alter when TSPs and ETPs transform into committed T-lineage precursors are largely unknown. Therefore, we inferred a trajectory from TSPs to TCR rearranging thymocytes (**Figure 4A**) through G1 annotated thymocytes (**Figure 2D**) along which differential gene expression analysis was performed. Consistently, this reflected the anticipated fluctuations in gene expression according to development, but not cell cycle (**Figure 4B**). Genes known to be expressed in multipotent hematopoietic progenitor cells (including *MEIS1, HOXA9, BCL11A, MEF2C* and *LMO2*) peak in the TSPs, as do the chemokine receptors *CCR7* and *CCR9*, consistent with their role in thymus homing in the mouse (Krueger et al., 2010; Zlotoff et al., 2010) (**Figure 4C**). Expression of *MME* (CD10) was sharply downregulated when human TSPs differentiate into ETPs under the influence of Notch signaling as evident from *DTX1* induction. During this Notch-dependent transition, the expression of the immature thymocyte markers *CD7*, *CD34* and *SELL* (CD62L) (Kohn et al., 2012) increases, consistent with observations when human HSCs are exposed to Notch ligands (Awong et al., 2009; Van de Walle et al., 2009). This coincides with a transient expression of other stem and progenitor genes such as *HHEX* and *LYL1*, and of *SPI1* that suggests multi-lineage priming as recently described during murine T cell development (Zhou et al., 2019). Differentiation into ETPs is also characterized by the upregulation of *GATA3* which continues to increase in expression during the subsequent specification and commitment stages (Van de Walle et al., 2016), but that clearly precedes expression of other critical T-lineage TFs such as *TCF12, TCF7* and *BCL11B* (Ha et al., 2017); and of the CD3 genes (*CD3D*, *CD3E* and *CD3G*) that all remain required for further T-lineage identity and function. Consistent with recent work, *BCL11B* induction preceded *CD1A* expression but was slightly delayed compared to loss of *CD44* expression, also at the protein level (**Figure 4D**), nevertheless confirming that the induction of T-lineage commitment occurs within the CD34^+^CD1a^-^thymocyte fraction (Canté-Barrett et al., 2017). Instead, *CD1A* expression seemingly marks the onset of TCR rearrangements given the similar induction kinetics of *PTCRA* expression and both RAG genes. This is closely followed by the induction of *CD1B* and *CD1E* (**Figure 4C**).

**Figure 4:**
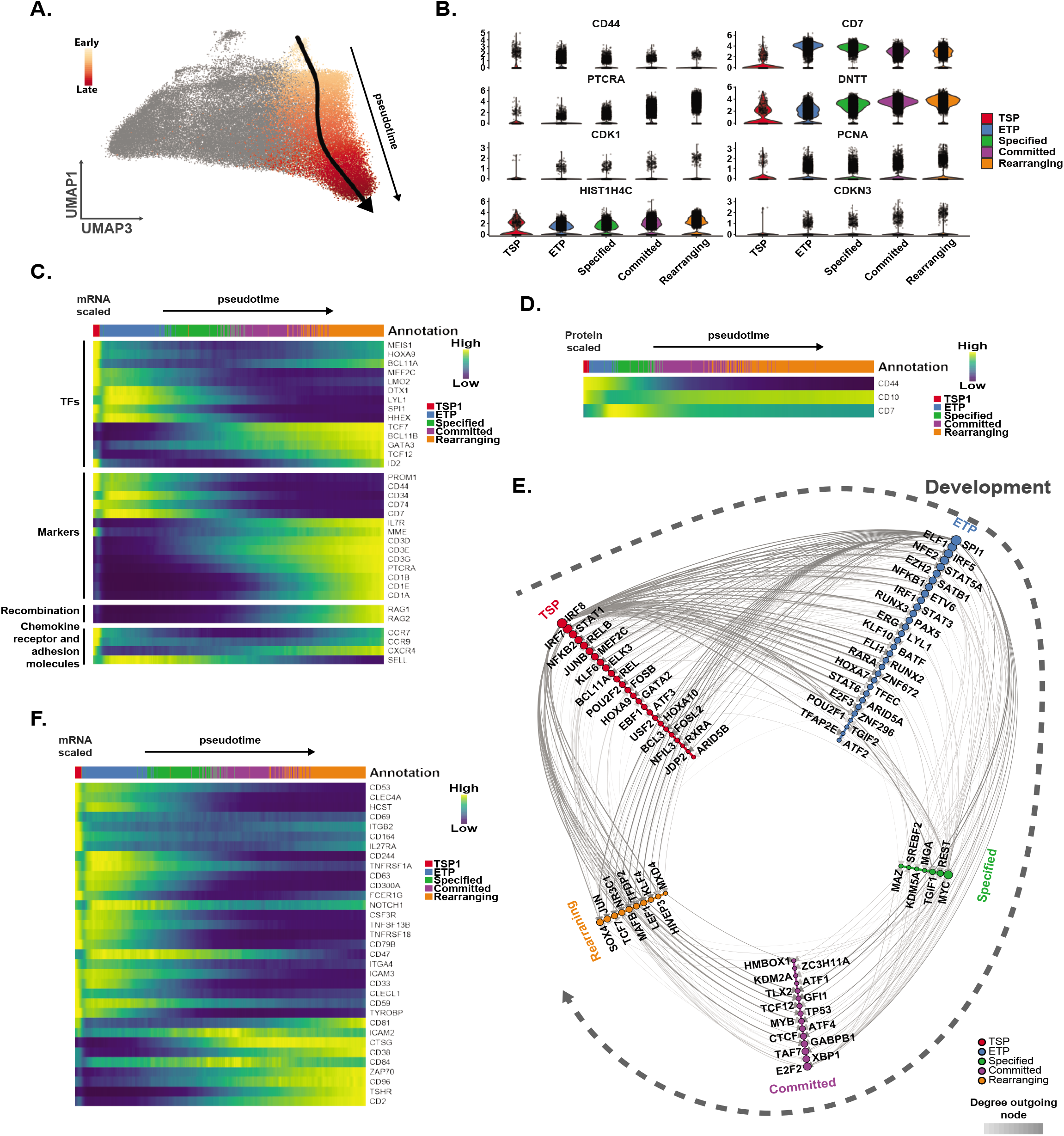
The transcriptional landscape of early human T lymphocyte development. **A.** UMAP visualization of the slingshot mediated trajectory inference across developing CD34^+^ G1 thymocytes. **B.** Violin plots visualizing expression of genes associated with early human T cell development and cell cycle across the annotated developing G1 thymocytes. **C.** Heatmap showing differential gene expression along the early CD34^+^ T cell differentiation pseudotime from TSP to rearranging thymocytes. Gene expression was visualized as a rolling average along pseudotime of DCA imputed log2 counts. **D.** Heatmap showing CITEseq-derived scaled protein expression data for CD44, CD10 and CD7 along the early T-lineage differentiation pseudotime. **E.** Hive plot showing the predicted network of transcriptional regulation among the differentially expressed transcription factors along the early T cell differentiation pseudotime. Regulatory interactions were predicted using the SCENIC algorithm and each axis corresponds to a developmental stage along the pseudotime. Transcription factor nodes across the axis were based on the maximal expression of each gene. Ordering along each axis is based on the amount of connections. Node size is dependent on the amount of connections, while edge size and opacity is based on the weight given by SCENIC. **F.** Heatmap showing differential expression of genes encoding for cell surface markers that are SCENIC predicted transcriptional targets of the transcriptional network depicted in E and ordered along the early T cell differentiation pseudotime. Gene expression was visualized as a rolling average along pseudotime of DCA imputed log2 counts.

To further improve our understanding of the TFs that drive early human T cell development, we implemented SCENIC (Aibar et al., 2017) which combines gene correlations with binding motif analysis within individual cells to infer regulons that correspond with co-expressed genes (**Figure 4E**). While this confirmed our TF gene expression patterns along pseudotime (**Figure S4A**), it created a more comprehensive overview of the transcriptional landscape that drives early human T cell development. Besides established stem and progenitor TFs (*MEF2C, BCL11A, GATA2* and *HOXA9*) (Yui and Rothenberg, 2014), SCENIC also revealed some activity of more specific, non T-lineage TFs (*IRF8, IRF7, EBF1, NFIL3*) in TSPs, thereby underscoring their multipotent properties. Notch-driven differentiation towards ETPs results in loss of activity of some of these early hematopoietic genes and induced IL7-dependent proliferation (*STAT5A, EZH2* and *E2F3*) Similarly to murine ETPs, *SPI1* (encoding for PU.1) displayed abundant transcriptional activity in these cells and was predicted to promote proliferation by activating *MYC*, *EZH2* and *E2F2*, as well as to maintain a multipotent transcriptional landscape by promoting the expression of genes associated with stem and progenitor cells (*MEF2C, BCL11A* and *HOXA9*). Specification towards the T-lineage apparently resulted in a less complex transcriptional landscape and revealed downregulation of genes associated with multipotency (*SPI1, LMO2* and *LYL1*), consistent with the requirement for cells to become committed to the T-cell fate. This process, coincided with a predicted peak in E-protein activity (*TCF12*) that is important for inducing *RAG* gene expression to permit TCR rearrangements (**Figure 4E**). These recombination events are further supported through epigenetic mechanisms (*CTCF*, *KDM2A*) and suppression of proliferation (*TP53*) and transcription (*GFI1*) (**Figure 4E**). Similarly, we employed SCENIC to retain predicted interactions to identify novel cell surface markers that marked particular stages of early human T cell development (**Figure S4B**). This validated that MME (CD10) and CD44 are initially both characteristic for TSPs but that they have opposing patterns during T-lineage commitment (**Figure 4C**), which was also confirmed at the protein level (**Figure 4D**). Additionally, we identified novel candidate cell surface markers for TSPs (*CD53* and *CD164*), for phenotyping the T-lineage specification stage (*HCST, CD69* and *CD244*) or to determine commitment (*CTSG*) (**Figure 4F**). Thus, these findings unravel the precise transcriptional dynamics through which TSPs are converted into committed T-lineage progenitors and proposes a gradual reduction in transcriptional network complexity as this process unfolds.

### Characterization of a primed thymus seeding progenitor population

Although previous studies also proposed that CD10 expressing lin^-^CD34^+^CD7^-^ thymocytes comprise the most immature thymus seeding precursors in human, others suggested that these precursors lack thymus homing potential and that instead CD34^+^CD7^int^ progenitors repopulate the human thymus (Haddad et al., 2006; Hao et al., 2008; Six et al., 2007) (**Figure 5A**). To further study this, we examined immature CD34^+^ peripheral blood mononuclear cells (PBMCs) and indeed observed a considerable fraction of CD34^+^CD7^int^ PBMCs (**Figure 5B**). Co-culture experiments on OP9-DLL1 stromal cells revealed that both CD34^+^CD7^-^ and CD34^+^CD7^int^ PBMCs progenitor populations display T-lineage potential and differentiate into committed CD7^+^CD1a^+^ thymocytes (**Figure 5C**). In addition, we observed no differences in thymus colonization between both populations using murine fetal thymic lobes in a hanging drop system (**Figure 5D**), in contrast to previous work that used the same approach (Haddad et al., 2006). Moreover, pertussis toxin (PTX) mediated blocking of G protein-coupled receptors, which include the thymus homing-associated chemokine receptors CCR7 and CCR9, demonstrated that both populations migrate into the thymic lobes through such specific mechanisms (**Figure 5D**).

**Figure 5:**
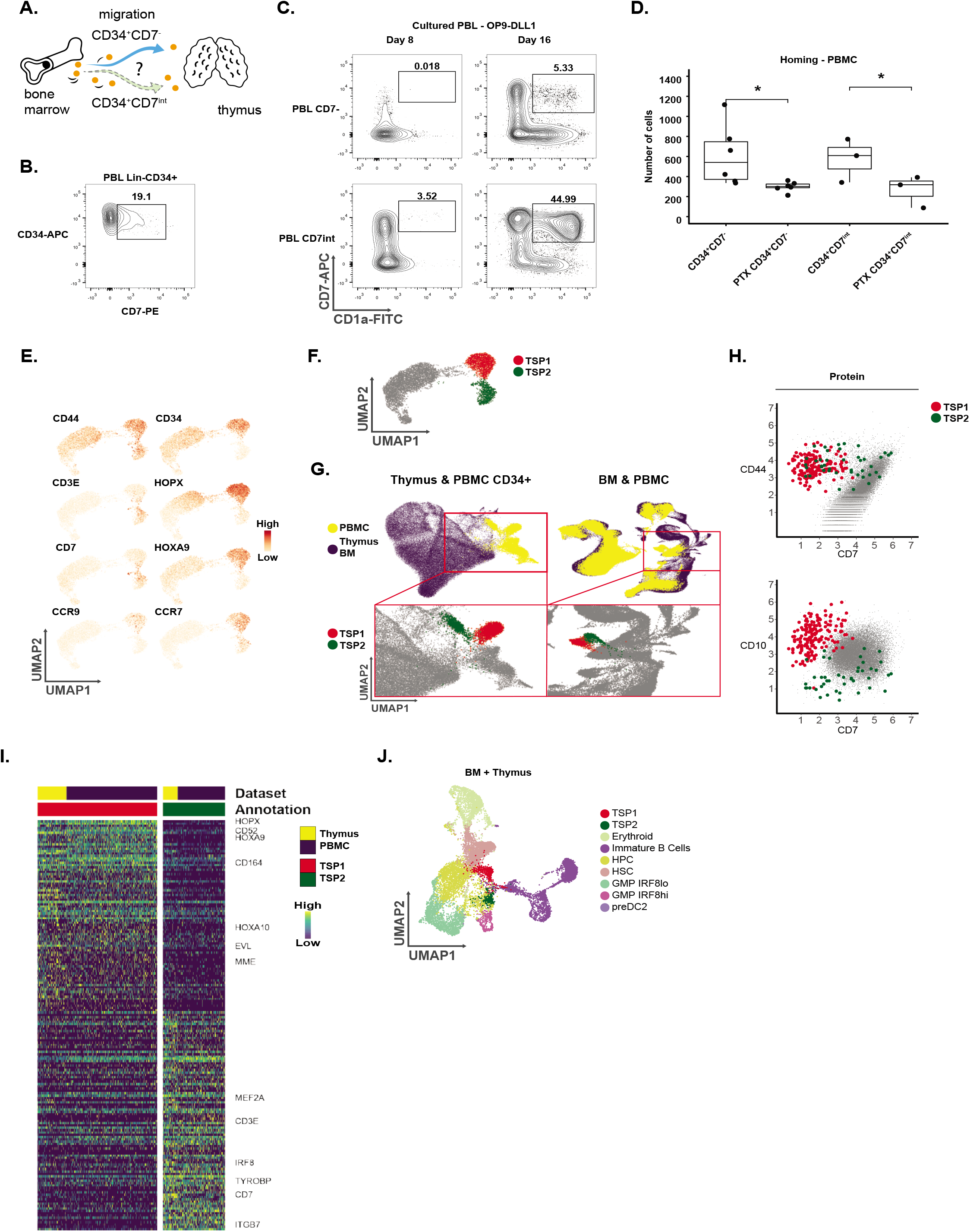
Functional assessment of human thymus seeding populations. **A.** Schematic overview of the migration of the newly identified lin^-^CD34^+^CD7^-^ putative TSP from the BM into the thymus in relation to the previously proposed lin^-^CD34^+^CD7^int^ TSPs (Haddad et al., 2006). **B.** Contour plot showing representative CD34 versus CD7 flow cytometric analysis of lin^-^CD34^+^ PBMC. Number in the density plot displays the frequency of CD7^int^ cells. **C.** Flow cytometric analysis of *ex vivo* isolated lin^-^CD34^+^CD7^-^ and lin^-^CD34^+^CD7^int^ PBMC cultured on OP9-DLL1 stromal cell during 8 and 16 days. Contour plots show CD7 versus CD1a staining to reveal T-lineage potential. Numbers in the density plots display the frequencies of CD7^+^CD1a^+^ T-lineage committed thymocytes. Data is representative for 3 independent experiments. **D.** Thymus seeding potential of lin^-^CD34^+^CD7^-^ and lin^-^CD34^+^CD7^int^ PBMCs following 48 hours in hanging drops in the absence of exogenous cytokines and in the absence or presence of pertussis toxin (PTX) to block G-coupled receptors. Box plots show data from 3-5 independent experiments (* p-value < 0.05, paired Student t-test). **E**. UMAP representing the expression of TSP specific and early T-lineage genes in the isolated population of immature PBMCs, thereby revealing a *CD7* and *CD3E* expressing subset that is distinct from the previously defined TSP subset. The size of dots corresponds to gene expression levels. **F.** UMAP visualization of the annotated putative TSP subsets in the immature PBMCs. **G.** UMAP visualization showing the overlay of the integration of either the immature PBMC fraction or entire PBMC dataset into the immature thymocyte (left) or the HCA BM dataset (right), respectively. Upper UMAP visualizes the overlay of the PBMC (yellow) and thymocyte (purple on the left) or BM (purple on the right) datasets. Lower UMAP visualizes both TSP populations in the overlap between the PBMC and thymocyte dataset (left) or the PBMC and HCA BM dataset (right). **H.** Scatterplots of protein surface expression of CD44 and CD7, and CD10 and CD7 through CITEseq (sort 2) for both populations of TSPs. The size of dots corresponds to gene expression levels. **I.** Heatmap visualizing the top 150 most differentially expressed genes between both TSP subsets. **J.** UMAP visualization of the annotated CD34^+^ BM cells and overlapping CD34^+^ G1 thymocyte populations following integration as shown in **Figure 2C** and updated to include the TSP2 population.

Next, we analyzed the published PBMC scRNAseq (10xGenomics; Zheng et al., 2017) dataset to search for migrating hematopoietic precursor cells that contain a transcriptional profile that matches TSPs. Within the cluster of *CD34* expressing cells (**Figure S5A**), we identified an isolated population with *CD44, CD34, HOPX, CCR9* and *CCR7* expression, transcriptionally corresponding to our proposed BM and thymus resident TSP population (**Figure 5E**). We also discovered a population of immature precursors lacking *HOPX, HOXA9, CCR9* and *CD34* expression, but expressing *CD44* and *CCR7*, as well as the Notch target genes *CD3E* and *CD7*, possibly reflecting the human counterpart of Notch-primed TSPs as described in mice (Chen et al., 2019; Yu et al., 2015). Therefore, we will refer to the initially identified population of immigrating progenitor cells as TSP1 and to the newly identified and presumably Notch-primed population as TSP2 (**Figure 5F**).

If TSP2 cells have the potential to migrate from the BM to seed the thymus, the transcriptional profile of the PBMC-derived TSP2 population should match with cells present in BM and thymus datasets. Indeed, we observed overlap for the TSP2 population with their thymic (**Figure 5G**, left) and BM-resident (**Figure 5G**, right) counterparts. Based on its topology, the TSP2 subset is transcriptionally distinct from the TSP1 population (**Figure 5F-G**) which was confirmed by differential gene expression analysis that revealed expression of *MEF2A, CD3E, CD7, IRF8, TYROBP* and *ITGB7* in TSP2, but not TSP1. This difference was preserved across the thymocytes and PBMCs (**Figure 5I**). Phenotypically, the TSP2 population differed from the TSP1 subset by lacking CD10 expression while predominantly being positive for CD7 (**Figure 5H**). Thus, these findings suggest that, within the pool of CD34^+^ BM-resident hematopoietic progenitors (**Figure 5J**), two distinct precursor subsets can be identified that differ in CD7 and CD10 surface markers that both can migrate through the blood and colonize the thymus.

### Dissecting the heterogeneity of the human thymic CD34+ fraction

Human T cell precursors have the potential to differentiate into multiple hematopoietic cell types when cultured *in vitro*, including erythroid, myeloid and lymphoid lineages, but whether this occurs *in vivo* has remained unclear (Hao et al., 2008; Márquez et al., 1995; Weerkamp et al., 2006). In addition, this apparent multilineage capacity may also result from heterogeneous precursor subsets that are present within the isolated phenotypes, a feature that was also identified in the most immature mouse DN1 thymocytes (Porritt et al., 2004). To identify the cellular heterogeneity within the CD34^+^ thymocytes, and to potentially reveal intrathymic non T-cell lineage differentiation, we integrated the HCA BM dataset, which should include these other differentiation pathways from the HSC stage onwards (**Figure 6A**), with our *ex vivo* immature thymocyte dataset with the exclusion of samples obtained from sort 3 (**Figure 1A**) since these contain mature lineage positive cells. Using this approach, we identified rare populations of erythroid and B-lineage progenitors within the CD34^+^ thymocytes, and most abundantly myeloid progenitors, including GMP populations and smaller but distinct DC precursor subsets (**Figure 6B-C**). We confirmed the identity of these cells using differential gene expression (**Figure 6D**). Inclusion of the sort 3 samples (**Figure S5B-C**) not only confirmed the consistent detection of these progenitor cells, but also revealed populations that transcriptionally corresponded to mature DCs (cDC1, cDC2 and pDC), mature B cells, plasma cells, monocytes and few mature NK cells (**Figure S5D-E**). After cell cycle regression, we could confirm the previously identified topological separation between TSP1 and TSP2, and detected multiple branches that particularly recapitulated myeloid development (**Figure 6E and Figure S6A**). This included a dominant pDC branch through a GMP IRF8hi precursor population as well as a cDC branch with specific precursors for both cDC1s and cDC2s and their progeny (**Figure 6E and S6B**). We did not find clear evidence for a NK/ILC precursors within the CD34^+^ thymocytes and the few mature NK cells that we identified were closest in topology with the TSP2 subset, not with ETPs or T-lineage specified precursors (**Figure 6E**).

**Figure 6:**
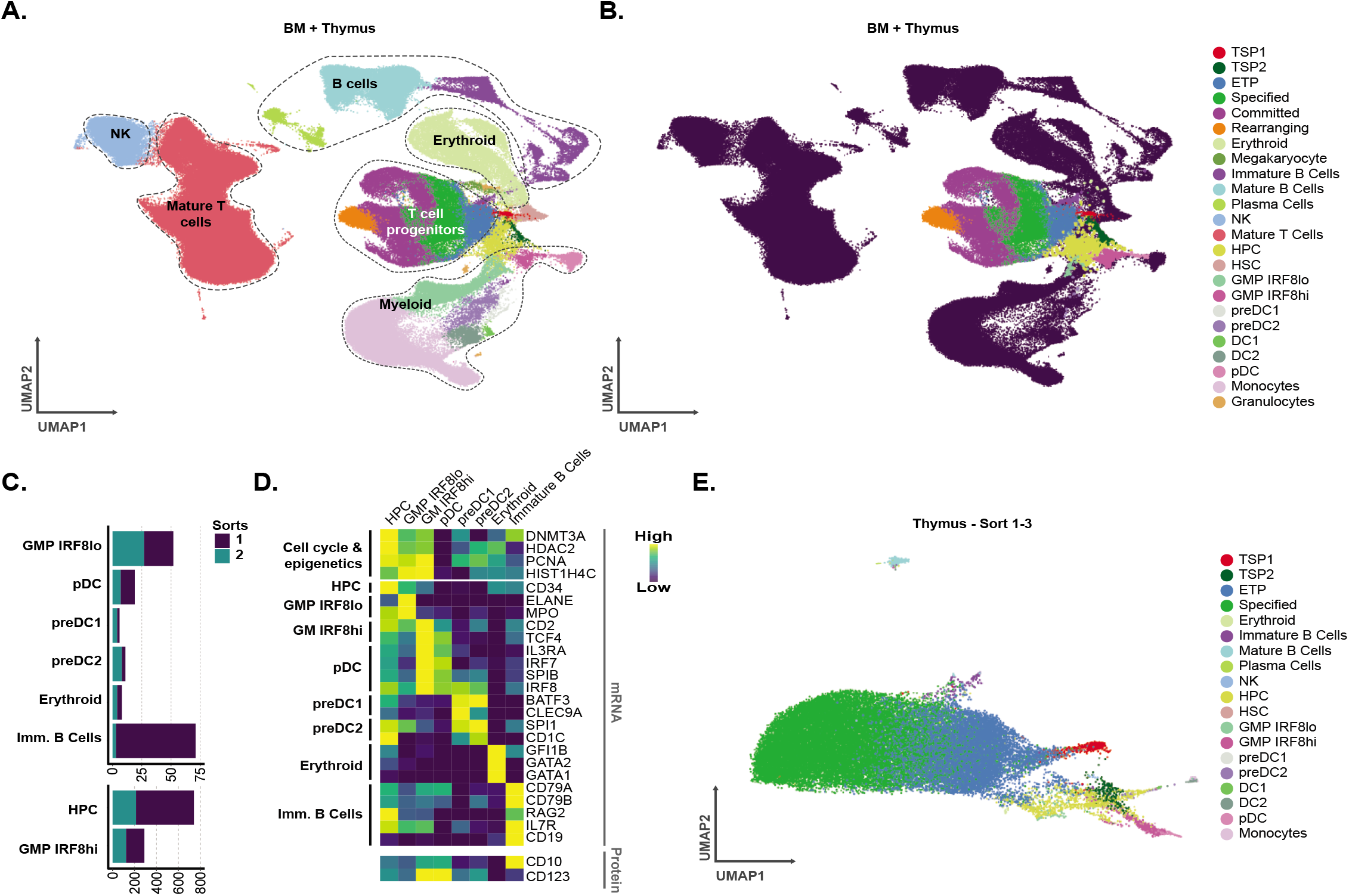
The CD34+ fraction of human thymocytes consists of a transcriptionally heterogenous group of cells. **A.** Annotated UMAP of the integrated *ex vivo* sorted human thymocytes (sort 1-2) and the HCA BM dataset. Error bars represent the standard deviation. **B.** UMAP visualization of the annotated subpopulations identified in the CD34^+^ human thymocytes (sort 1-2). **C.** Stacked bar chart visualizing the amount of detected populations for sort 1-2. **D.** Heatmap visualizing scaled and averaged mRNA and protein expression characteristic for the identified sub populations (n > 3) (Sort 1-2). **E.** Annotation of developmental subsets (n > 3) visualized on a UMAP of the uncommitted CD34^+^ fraction following cell cycle regression (Sort 1-3).

To clarify whether the identified thymic non T-lineage progenitor cells develop intrathymically, or instead migrate from the periphery into the thymus, we annotated the immature fraction of PBMCs (Figure 7A) which should contain all putative migratory subsets. The TSPs displayed an increased frequency in the periphery compared to the thymus (Figure 7B) and predominantly consisted out of G1 cells (Figure 7C). Also intrathymic immature B cells consisted primarily of G1 cells. In contrast, GMP and HPC populations were found to be proliferative in a thymic environment (Figure 7C) and less abundant in the periphery (Figure 7B). The hypothesis that TSP1, TSP2 and immature B cells migrate into the thymus was further supported by integrating the *ex vivo* human thymocyte, CD34^+^ BM and immature PBMC datasets which revealed overlap in these tissues (Figure S7A-B). Importantly, we could not detect any HSCs in our thymus dataset while these cells were abundant in the BM and PBMCs, indicating that our thymus-derived cells were not contaminated with blood cells. Within the thymus, the GMP and HPC populations display evidence of Notch signaling as the majority of cells expressed CD7 protein (Figure 7D), suggesting intrathymic expansion and development out of a blood-derived migratory population. In line with these findings, the thymus was found to harbour all DC intermediates while GMP IRF8hi precursors that branch towards pDCs were not detected in the blood (Figure 7E-F). Overall, these findings suggest that multiple progenitor cell types can migrate into the thymus. While some do not alter their properties in response to the thymic microenvironment, others, like HPCs, proliferate upon Notch activation and presumably support T cell development by serving as a precursor source for both T- and non T-cell lineages. Particularly differentiation towards pDCs seems to be supported by the thymus through the generation of an intrathymic GMP IRF8hi progenitor population. In contrast, clear signs of intrathymic NK cell development were missing.

**Figure 7:**
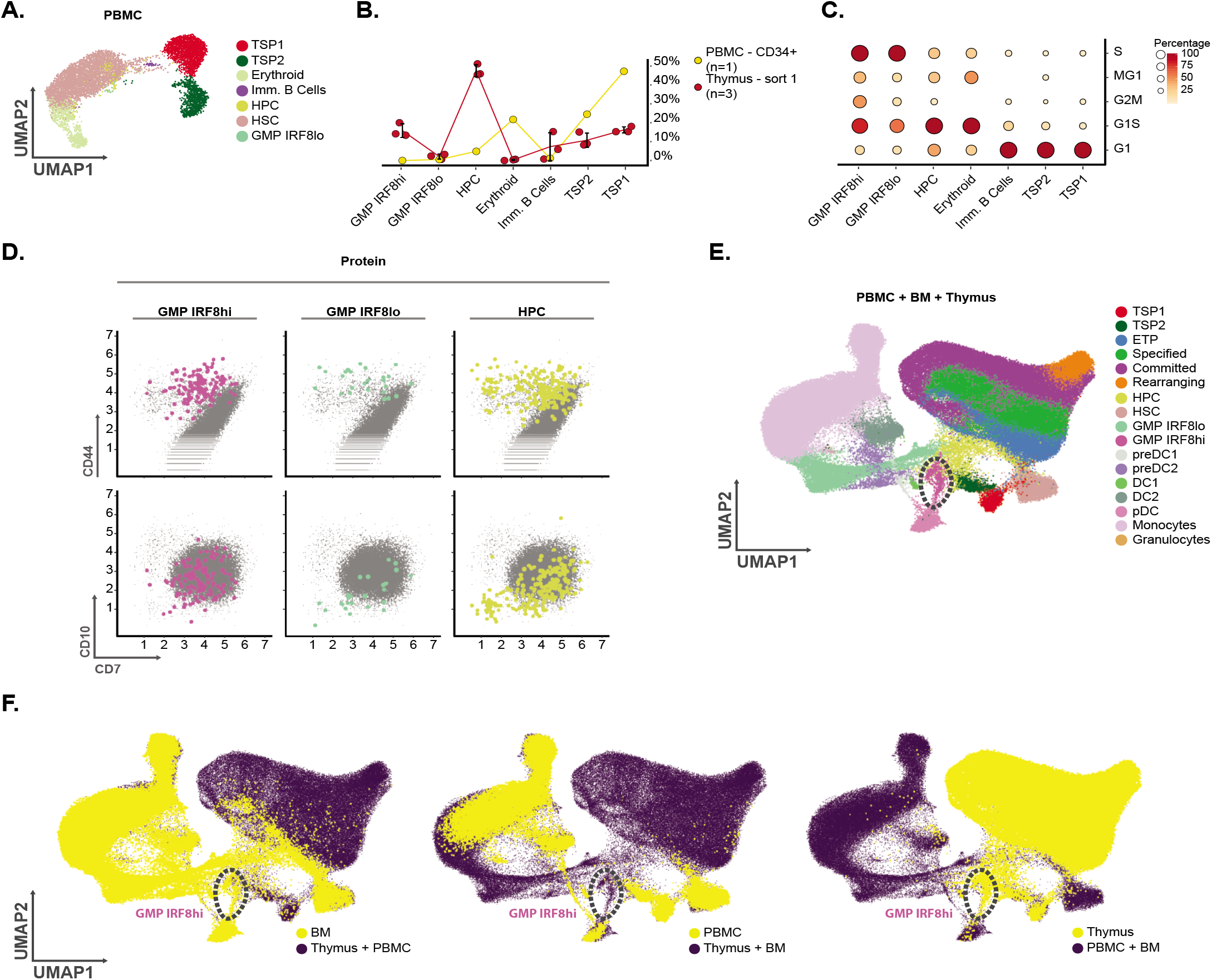
Intrathymic development of plasmacytoid dendritic cells. **A.** UMAP visualization of the annotated immature PBMC cells. **B.** Comparison between the frequency of subsets in the lin^-^CD34^+^CD1a^-^ thymocytes (sort 1) and CD34^+^ PBMCs. **C.** Dot plots visualizing the cellular frequency for each developmental subset according to specific cycle stages. **D.** Scatterplots of protein surface expression of CD44 and CD7, and CD10 and CD7 through CITEseq (sort 2) for the most abundant CD34^+^ thymocyte populations. **E.** Annotated UMAP of the integrated BM cells, PBMCs and *ex vivo* thymocytes (sort 1-3), with the GMP IRF8hi cells marked. **F.** Overlays of the integrated datasets as described in E where either the BM (left), PBMC (middle), *ex vivo t*hymocyte (right) are highlighted in yellow, while the remainder of cells are displayed in purple. The GMP IRF8hi cells, absent in PBMCs, are marked

## Discussion

The lack of genetic tools and the absence of comparable cell surface markers between mouse and human have impeded our insights into the early stages of intrathymic T cell development in human. Here, we profiled approximately 70.000 individual CD34^+^ pediatric thymocytes through scRNAseq to obtain a high-resolution cellular atlas of the early stages of human intrathymic hematopoiesis. Integration with BM and PBMC data allowed us to identify various precursor subsets that seed the thymus, and to delineate intrathymic differentiation towards T- and non T-cell lineages. By including CITEseq, we were able to link our annotated precursor populations to various phenotypically identified human T cell precursors in literature, thereby resolving these seemingly conflicting studies. This study thus clarifies the heterogeneity of early T cell precursors in human and provides clear insights into the precursor cells and their transcriptional dynamics that drive human T cell development after birth. This resource will therefore serve as important framework for optimizing T cell reconstitution in T cell deficient patients.

During postnatal life, thymopoiesis is maintained by the colonization of BM-derived progenitors of which only few can seed the thymus, presumably only upon niche availability. In human, multiple populations have been proposed to represent these immature thymocytes based on differential CD7, CD10 or CD38 expression and *in vitro* differentiation experiments were used to investigate their hematopoietic lineage potential (Dik et al., 2005; Haddad et al., 2006; Hao et al., 2008; Six et al., 2007). A major strength of our study, to resolve the nature of the thymus seeding precursor, is that we not only performed scRNAseq on a large number of immature thymocytes, but that we also implemented CITEseq to link the transcriptome with phenotypic properties and used integration with other datasets to demonstrate their migratory potential. Using these approaches, we identified two non-proliferative populations which are present in the thymus, BM and blood. Of these, the TSP1 transcriptionally corresponds to murine TSPs based on expression of chemokine receptors (*CCR7* and *CCR9*) and transcription factors (*HOXA9, MEIS1* and *MEF2C/* (Yui and Rothenberg, 2014). Immunophenotypically, TSP1s are defined as Lin^-^CD34^+^CD44^hi^CD7^-^CD10^+^, consistent with the studies from the Crooks and Andre-Schmutz labs (Hao et al., 2008; Six et al., 2007) and the initial work from Galy et al (Galy et al., 1995). Although previously thought to lack the potential to colonize the thymus, we could demonstrate that CD7^-^ precursors can actively colonize thymic lobes through the activity of G protein-coupled receptors, consistent with the expression of *CCR7* and *CCR9* in these precursors. These findings are also consistent with the first studies that used chimeric FTOCs with human hematopoietic progenitors since CD7^+^ HPCs were excluded and considered lineage positive cells at that time (Plum et al., 1999; Taghon et al., 2002). Also their topology within the thymus suggests that these TSP1s represent the canonical T cell precursors that differentiate into ETPs when they encounter Notch activating signals within the thymic microenvironment. And while TSP1s themselves do not display any signs of Notch activation, we also identified a lin^-^CD34^+^CD44^hi^CD7^int^CD10^-^ seeding population which resembles the Notch-primed TSPs that have been described in the BM of mice (Chen et al., 2019; Yu et al., 2015). This TSP2 population indeed expresses the Notch target genes *CD7* and *CD3E* prior to thymic entry, consistent with these studies. Importantly, both TSP1 and TSP2 are subfractions of the lin^-^CD34^+^CD7^-^ and lin^-^CD34^+^CD7^int^ thymocytes that were previously sorted and differentiated in vitro to test their lineage differentiation potential. Our study reveals that these populations thus harboured multiple precursor subsets that will have influenced these in vitro differentiation experiments and that only a fraction of these cells were actual TSPs. Nevertheless, our work nuances previous findings by suggesting that at least two distinct populations can seed the thymus, thereby also reconciling these studies. Importantly, these TSP subpopulations differ from the recently described human fetal TSPs (Zeng et al., 2019) which where shown to express high levels of *IL3RA*, *IRF8* and *IL6R*. The unique transcriptional profile of the postnatal TSPs will clearly be important from a translational perspective to improve T cell reconstitution in patients following for instance hematopoietic stem cell transplantations.

Using TSP1 as the canonical T cell progenitor, we inferred a trajectory to reveal the transcriptional changes during early T cell development in human and employed SCENIC to predict the transcriptional networks that control the discrete steps of this process. Consistently, TSP1 cells expressed high levels of stem cell and progenitor genes which are gradually downregulated during the ETP stage, consistent with recent data from murine T cell development (Zhou et al., 2019). However, these TSP1s also display transcriptional activity of more lineage specific transcription factors such as EBF1, IRF8 and NFIL3, which was not as obvious in the most immature mouse thymocytes. Such multilineage priming is also evident in human ETPs as shown through *SPI1* and *MPO* expression, similar as in mouse, and consistent with the lineage potential of these immature cells when extracted from the thymus (Hao et al., 2008; Márquez et al., 1995; Six et al., 2007; Weerkamp et al., 2006). ETPs also displayed the first signs of IL7R signaling and expressed the highest levels of the Notch target genes DTX1 and CD7. Such findings are consistent with earlier studies from our lab that showed that human ETP generation requires the strongest levels of Notch activation during T cell development and that further T-lineage commitment and differentiation requires downmodulation of Notch activity, a process that is dependent on GATA3 activity (Van de Walle et al., 2016). Our single cell data confirms this switch from Notch to GATA3 activity and this process also coincides with the downregulation of CD44, recently described as being a Notch target gene (García-Peydró et al., 2018) whose downregulation matches human T-lineage commitment (Canté-Barrett et al., 2017). BCL11B has also been shown to regulate human T-lineage commitment but its induction comes later compared to GATA3. This may suggest that GATA3 initiates T-lineage commitment while BCL11B rather enforces it (Ha et al., 2017; Li et al., 2013). Strikingly, induction of TCF7 expression coincides with BCL11B induction which is much later compared to during murine T cell development (Weber et al., 2011), indicating that the transcription factor networks that control these distinct stages of human T cell development are different compared to in mouse. Thus, our work not only strengthens previously reported findings regarding either the immunophenotypic changes or transcriptional drivers at play during early human T cell development, but also builds upon these findings by identifying novel and stage-specific transcriptional networks. Together with the potential newly identified cell surface markers that seemingly characterize specific precursor subsets or stages of human T cell development, this study therefore also provides novel avenues for further exploration and validation to fully unravel the complexity of early T cell development in human.

Our work also provides interesting novel hints with respect to the non T-cell lineages that actually develop in vivo within the thymic microenvironment, as issue that has been complicated due to the lack of genetic tools to study human thymopoiesis in vivo. Our results indicate that B and erythroid lineages do not develop in the human postnatal thymus, but rather originate from lineage-specific precursors that immigrate from the blood. This may also explain why it has been difficult to reveal consistent B and erythroid potential from intrathymic progenitors (Hao et al., 2008; Weerkamp et al., 2006). From our scRNAseq, we could only find convincing evidence for the in vivo intrathymic development of DC-lineage cells, possibly from immigrating HPCs that expand in response to Notch signaling, consistent with recent studies (Kirkling et al., 2018). Particularly pDC development was evident through an IRF8^hi^ expressing GMP precursor that most likely develops in the thymus given their virtual absence in the blood. These precursors may also correspond to the recently described CD34^+^CD123^+^ precursors that display DC potential (Martín-Gayo et al., 2017). Thus, these findings reveal that the thymus supports DC development, but presumably from a distinct precursor compared to the thymocytes that we believe to mainly originate from TSP1s. Strikingly, we could find virtually no evidence for NK cell development in the postnatal thymus. Very few NK cells were detected, and these were transcriptionally most similar to the TSP2 subset. We certainly did not see any branch of developing NK cells from ETPs or T-lineage specified cells, suggesting that the residual NK-lineage potential in these cells results from the similar transcriptional networks that drive their development (De Obaldia and Bhandoola, 2015), rather than being an actual event that occurs in the thymus. While the lack of NK cell development is surprising, this finding is however consistent with observations in SCID patients that reconstituted T cells but failed to develop NK cells following HSC transplantation, without adverse effects (Vély et al., 2016).

In summary, we have generated a unique and high-resolution dataset which yields detailed information about both the cellular diversity of immigrating and developing human CD34^+^ thymocytes, as well as their underlying transcriptional landscape and immunophenotype. This provides innovative insights into the intrathymic lineage differentiation events in human and the supporting transcriptional networks that await further exploration.

## Material and methods

### Isolation of human CD34+ hematopoietic progenitor cells from peripheral blood and postnatal thymus

Peripheral blood from healthy donors and pediatric thymus from children undergoing cardiac surgery were obtained and used according to and with the approval of the Medical Ethical Commission of Ghent University Hospital (Belgium). Peripheral blood mononuclear cells (PBMCs) were isolated from peripheral blood following Lymphoprep density gradient while the thymus tissue was mechanically disrupted to obtain a single cell suspension. CD34^+^ PBL and CD34^+^ thymocytes were enriched using MACS with CD34 microbeads (Miltenyi Biotec) according to the instructions of the manufacturer. PBL cells were sorted as CD34^+^CD19^-^CD14^-^CD3^-^CD56^-^(Lin^-^)CD7^-^and CD34^+^Lin^-^CD7^+^ cells for OP9-DLL1 coculture and thymus homing experiments. Enriched CD34^+^ thymocytes were sorted into the lin-CD34+CD1a- (Sort 1), lin^-^CD34^+^CD7^-/int^, lin^-^CD34^+^CD1a^-^ and lin^-^CD34^+^CD1a^+^ (sort 2), lin^+^, lin^-^ CD44^+^CD7^-/+^ and lin^-^CD44^int^CD7^+^ (sort 3) populations for scRNAseq; and into CD34^+^CD1a^-^CD7^-^ and CD34^+^CD1a^-^CD7^+^ fractions for OP9-coculture experiments.

### Thymus homing assay

CD34^+^Lin^-^CD7^-^ and CD34^+^Lin^-^CD7^+^ PBMCs were incubated for 2 hours at 37°C and 7% CO2 in complete IMDM in the absence or presence of 100 ng/ml of pertussis toxin to block the activity of G-coupled chemokine receptors. Subsequently, the cells were washed twice and incubated with thymic lobes from fetal day 14-15 NOD-SCID mice in a Terasaki plate at a density of 25.000 cells per lobe in a volume of 25 μl of complete IMDM. The Terasaki plates were inverted to form hanging drops during 48 hours before the lobes were isolated and rigorously washed in PBS to remove non-colonized cells. Thymic lobes were mechanically disrupted with a tissue grinder to obtain a single cell suspension that was used for quantification of human cells by flow cytometry analysis using BD counting and human CD45 labeling.

### Co-culture experiments for scRNAseq or flow cytometric analysis

To investigate T-lineage potential, thymus or PBMC derived CD34+ precursor subsets were cultured on OP9-DLL1 stromal cells in complete MEMa medium supplemented with 5 ng/ml IL7, 5ng/ml SCF and 5ng/ml SCF.

### Library preparation and sequencing

The sorted lin^-^CD34^+^CD7^-/int^, lin^-^CD34^+^CD1a^-^ and lin^-^CD34^+^CD1a^+^ thymocyte populations were pooled, using all available lin^-^CD34^+^CD7^-/int^ and equal amounts of the lin^-^CD34^+^CD1a^-^ and lin^-^ CD34^+^CD1a^+^ populations (sort 2). In addition, the sorted lin^+^, lin^-^CD44^+^CD7^-/+^ and lin^-^CD44^int^CD7^+^ fractions were pooled using all lin^+^ cells and a 5/3 ratio of the lin^-^CD44^+^CD7^-/+^ and lin^-^CD44^int^CD7^+^ populations (sort 3).

TotalSeq antibodies were added in concentrations as recommended by the manufacturer and stained for 30 min on ice. Finally, scRNAseq and CITEseq libraries were generated according to the instructions of the manufacturer.

### Preprocessing of the scRNAseq and CITEseq data

Using CellRanger 1.2.0, the sequencing data was demultiplexed and mapped against the GRCh38 genome. The resulting count matrices were loaded into R using the DropletUtils library. Additionally, the publicly available HCA BM dataset was downloaded from the HCA portal and loaded into R using Seurat (Butler et al., 2018).

Low quality cells were identified as having less than 500 genes, less than 1000 UMI and more than 10% mitochondrial reads. Using scrublet (Wolock et al., 2019) doublets were identified and both clusters containing doublets and outlier cells within the clusters were removed. Low quality genes were identified as being expressed in less than 3 cells. Additionally, genes encoded by either the Y chromosome or mitochondrial genome were removed as well. Subsequently, the high-quality cells were normalized using Scran (Lun et al., 2016a, 2016b).

The ADT libraries were mapped using the Cite-seq count program (Roelli et al., 2019). Following mapping, only the cells which were detected in both the scRNAseq and ADT libraries were retained in the ADT matrices. Subsequently, the Seurat library was used to normalized these count matrices.

### Highly variable gene selection, batch correction, dimensionality reduction and clustering

Using scran, highly variable genes (HVG) were identified by fitting a trend through the variances of the normalized log transformed counts and an appropriate design matrix was provided, taking information about the libraries, donors and gender into account, if applicable. Genes with a FDR smaller than or equal to 0.01 were considered significant and the top 4000 genes with the highest biological components, as determined by scran, were retained. Subsequently, for each library (Thymus and PBMC) or donor (HCA BM) a pca analysis was performed, from which the first 50 components were retained. These matrices were supplied to the FastMNN algorithm (k = 20), based on mnncorrect (Haghverdi et al., 2018), and batch effects were corrected, resulting in a corrected pca matrix. This batch effect corrected pca was used to generate an anndata object using the python package scanpy (Wolf et al., 2018), in R via reticulate. Using scanpy a knn graph (k = 15) was constructed and a UMAP (McInnes et al., 2018) manifold was generated. Leiden and Louvain clustering were done using scanpy, whereas walktrap and label propagation clustering were performed via the python igraph package.

### Dataset integration and batch correction

Datasets obtained from different tissues, such as the thymus, peripheral blood and bone marrow, were normalized separately and thus treated as different datasets. In order to integrate these different datasets, the size factors were rescaled using the multiBatchNorm function to remove systematic differences in coverage. Subsequently, highly variable genes were selected for each dataset separately. The resulting p values for each HVG analysis were combined using the combinePValues function in scran, followed by multiple testing correction (FDR); and the average biological component was calculated for each gene. Subsequently, genes with an FDR less than or equal to 0.01 were retained and the top 4000 genes with the highest averaged biological component were selected. The datasets were batch corrected as described above.

### Annotation of cell types across multiple datasets

In order to distinguish rare subtypes present in our *ex vivo* human thymocyte dataset, we performed stepwise integration and annotation of several datasets. This was done by performing Leiden clustering and manually annotating these clusters based on marker genes. Subsequently, the annotation was extended to unannotated clusters by implementing logistic regression (solver = saga) from the sklearn python module, in R via reticulate. This was done on the top 4000 most HVG in order to improve performance. Training datasets were generated by randomly subsampling the annotated clusters up to a maximum of 1000 cells per cell type, if possible. This was done separately for the ex vivo thymus and HCA BM datasets.

The thymocyte annotation was validated and extended by integrating the *ex vivo* thymocyte and HCA BM dataset as described above. Following integration, the annotation provided to the BM dataset was superimposed onto the ex vivo thymocyte dataset by using logistic regression, as described above. While this allowed annotation of rare thymocyte populations, such as erythroid progenitors, this also resulted in incorrect annotation of thymocytes. Therefore, using the neighbourhood cleaning rule (NCR) method (k=10), from the sklearn python module, putative outliers for each subset were predicted based on their position in the UMAP manifold. Subsequently, mutual nearest neighbours (MNNs) between the putative outlier and correctly annotated cells were identified using the findMutualMNN function (k=100) from the batchelor library. This was done until all putative outlier cells were paired with at least 5 correctly annotated cells. Next, for each of these MNNs the euclidean distance was calculated using the pairwise_distances function in the sklearn module, in R via reticulate. The obtained distances between the MNN pairs were subsequently weighted by calculating the variance between the euclidean distances of a specific outlier and it’s correctly annotated neighbours, which correspond to a specific annotation. These variances were added up to the euclidean distances between an outlier and it’s neighbours corresponding to a specific annotation, resulting in weighted distances. The median weighted distance was calculated for every outlier which had neighbours that corresponded to a specific annotation.

Based on the smallest weighted distance the outlier was provided with an annotation. Finally, the annotations provided by either logistic regression or based on the weighted distance were manually checked and corrected using a cluster-based approach (Leiden, Louvain or label propagation). Finally, this annotation was checked and corrected by either subsetting or integration with other datasets, such as the cultured thymocytes or immature PBMC.

The robustness of the resulting annotation was tested by randomly subsampling each population to contain maximally 2500 cells, if possible. Using the normalized log transformed counts the euclidean distances between the cells were calculated with the pairwise_distances function from the sklearn module, and hierarchical clustering was performed using ward linkage. The resulting distance matrix was visualized using the seaborn module in python.

The immature PBMC were annotated by superimposing the annotation from the BM cells using logistic regression, as described above. Using clustering the annotation was corrected.However, in contrast to the HCA BM dataset, not as many incorrectly annotated cells were identified, which is why manual correction based on Leiden clustering sufficed.

### Annotation and regression of cell cycle stages in the human *ex vivo* thymocyte dataset

Highly variable (FDR <= 0.01) cell cycle stage specific genes (Macosko et al., 2015) were retained from the libraries sequenced to a depth of 50000 reads/cell (sort 1). Subsequently, for each library pca matrices were generated and batch correction was performed using the FastMNN function in scran on the first 25 components of each pca matrix. Next, scanpy was used on the corrected matrix to perform Leiden clustering. Genes specific to either the G1S, S, G2M, M or MG1 stages were averaged (feature agglomeration) for each cell and scaled, which in turn was used to guide cluster annotation.

The obtained annotation was extended using the logistic regression function (solver = saga) from the sklearn python module, which was run in R via reticulate. In order to provide the algorithm with an appropriate training dataset each cell cycle stage was randomly subsetted to contain maximally 1000 cells, while retaining all significant HVG (FDR <= 0.01).

In order to remove cell cycle effects from the dataset, the vector containing the cell cycle annotation was included in the design matrix during HVG selection. From the resulting top 4000 HVG the cell cycle effects were regressed using the vector of cell cycle stages, prior to batch correction using the FastMNN function.

### Differential expression analysis on subsets

To identify differentially expressed genes across various developmental subsets, the limma package (Ritchie et al., 2015) was used, based on a recent benchmark (Soneson and Robinson, 2018). Instead of log2 transformed counts, scater (McCarthy et al., 2017) was used to calculate log2 transformed counts per million (CPM) using size factors obtained during normalization with scran. Subsequently, a design matrix was generated to specify confounding variables, such as library, donor, gender and sequencing depth (UMI counts and number of genes), while a contrast matrix was generated to incorporate these variables in the following log ratio test (LRT). the multiple testing error was corrected using the FDR method and genes with a FDR smaller or equal to 0.05 were considered significant. In order to identify up or down regulated genes, log fold changes were subtracted from each other and varying cutoffs were implemented.

### Heatmap visualization of pseudobulk normalized scRNAseq

In order to apply the statistical rigor of bulk RNAseq to scRNAseq the counts for each gene were summed for each developmental subset and sequencing library, using the sumCountsAcrossCells function from the scater package, resulting in pseudo bulk counts (Tung et al., 2017). The resulting matrix was normalized using the DESeq2 package and log transformed. Subsequently, gene expression was averaged for the donors, scaled gene-wise using the scale_minmax function from the SCORPIUS library and visualized using the pheatmap library.

### Trajectory inference and differential gene expression analysis

Slingshot (Street et al., 2018) was selected based on a benchmarking study (Saelens et al., 2019) to infer a trajectory of developing G1 thymocytes, through the defined developmental stages, instead of clusters, of the 3 UMAP dimensions. The resulting trajectory was used to arrange G1 thymocytes, confounding factors, such as library, donor, sex and sequencing depth, were regressed from the log2 CPM values and the resulting residuals were used to perform differential gene expression using a generalized additive model (GAM) (Perraudeau et al., 2017). In order to correct for multiple testing, the FDR was calculated and genes with a FDR lower than or equal to 0.05 were considered significant. Differentially expressed genes obtained in this manner and visualized using a heatmap underwent quantile scaling (SCORPIUS package), a rolling average along the trajectory was calculated for each gene and the heatmap was visualized using pheatmap.

Alternatively, RNA velocyto (La Manno et al., 2018) was used to infer a developmental trajectory. More specifically, loom files for each library were generated using the command line velocyto tool (version 0.17.17). These loom files were read into R using scVelo (Bergen et al., 2019) via reticulate and combined with the AnnData object containing a count matrix and the batch corrected pca matrix obtained following FastMNN (Haghverdi et al., 2018). Using scanpy the count matrix was normalized and log transformed. In order to run scVelo a python environment was used, where the RNA velocyto algorithm was run in stochastic mode on 2 dimensions of a UMAP manifold.

### SCENIC and graphs

Transcription factor (TF) activity was predicted by SCENIC (Aibar et al., 2017), using the python implementation, pySCENIC, and GRNBoost 2 (Moerman et al., 2019). Duplicated genes and low-quality genes were removed from the normalized log transformed count matrix in R. Subsequently, GRNBoost2 was used to determine correlation between transcription factors and other genes. Transcription factors correlating with at least 20 genes were retained and regulons were inferred based on publicly available motif binding databases provided by the Aerts lab.

The resulting area under the curve (AUC) matrices were quantile scaled with the SCORPIUS package (Cannoodt et al., 2016), ordered along the trajectory obtained following TI and the pheatmap package was used to visualize positive regulons as a heatmap.

In order to visualize TF activity in a cell type specific manner graphs were generated. These graphs combine the output of DE analysis, the AUC matrix generated with pySCENIC and the TF motif binding predicted by pySCENIC; all of the aforementioned methods are described above. Here, we define TF activity as interactions between a TF and its target genes predicted by pySCENIC, which is based on a combination of correlation between a TF and at least 20 genes, and motif binding databases which report the TF to bind these highly correlated target genes.

From the motif binding output predicted by pySCENIC, TFs and their target TFs were retained if they were differentially expressed, according to the GAM. The motif binding output from pySCENIC, where the weight attributed to each predicted TF-TF interaction, was used to filter TFs, retaining interactions with a weight above the median. Additionally, the retained TFs needed to correspond to positive regulons or were removed otherwise and have at least 2 interacting partners after filtering. The resulting edgelist was used to generate a graph using the igraph R library, which was in turn visualized as a hive plot using the ggraph library. In order to provide an identity to each TF, the cells were ordered along pseudotime, quantile scaled and a loess curve (span = 0.5) was fitted through each gene, where the maximum of the loess curve corresponded to a specific cell type.

Finally, interesting and significantly expressed (FDR <= 0.05) genes which encoded for cell surface markers and were downstream targets of TFs visualized via a hive plot were retained from the motif binding output from pySCENIC. Subsequently, from the resulting edgelist a graph was generated using the igraph R library and ggraph was used to visualize the interactions as a bipartite graph.

### Software

Data analysis of both the scRNAseq or CITEseq data was performed using either R (version 3.6.0) or python (version >= 3.6.4). Statistical analysis of homing experiments was done using R. Analysis of multicolor flow cytometry or FACS data was done using FlowJo (version 10.0.7).

## Acknowledgments

We wish to thank Sophie Vermaut and Katia Reynvoet for flow cytometry help, Juan-Carlos Zuniga-Pflucker (University of Toronto) for OP9 stromal cells, Dr. Francois and Dr. Van Nooten (Department of Cardiac Surgery; University Hospital Ghent) for thymus tissue and the Red Cross Flanders for peripheral blood. This work was supported by the Fund for Scientific Research Flanders, the Concerted Research Action of Ghent University Research Fund (GOA), the Foundation against Cancer (STK), the European Research Council (StG-639784 to PVV), Stand up to Cancer (the Flemish cancer society), the Belgian Cancer Society and the Chan Zuckerberg Initiative (CZF2019-002445). J.-E.P. is supported by an EMBO Long-Term Fellowship, S.A.T. was funded by Wellcome (WT206194), ERC Consolidator (no. 646794) and EU MRG-Grammar awards.

## Author contributions

M.L. performed the sample preparation, bioinformatics analysis, designed the study and wrote the paper. K.L.L., J.P. helped with conceptualization of the study. N.V. performed single cell RNAseq. J.R. helped with writing of the manuscript. M.S.K, B.L., O.A., M.T., D.D., T.L.T, M.S., O.R-R., A.R. generated bone marrow single cell RNAseq datasets. B.V., G.L. provided expertise. P.V.V., M.G., Y.S.,S.A.T. assisted with funding acquisition and provided resources. T.T. Performed experiments, design and supervision of the study, acquisition of funding, wrote the paper.

## Supplementary Figure Legends

**Figure S1:**
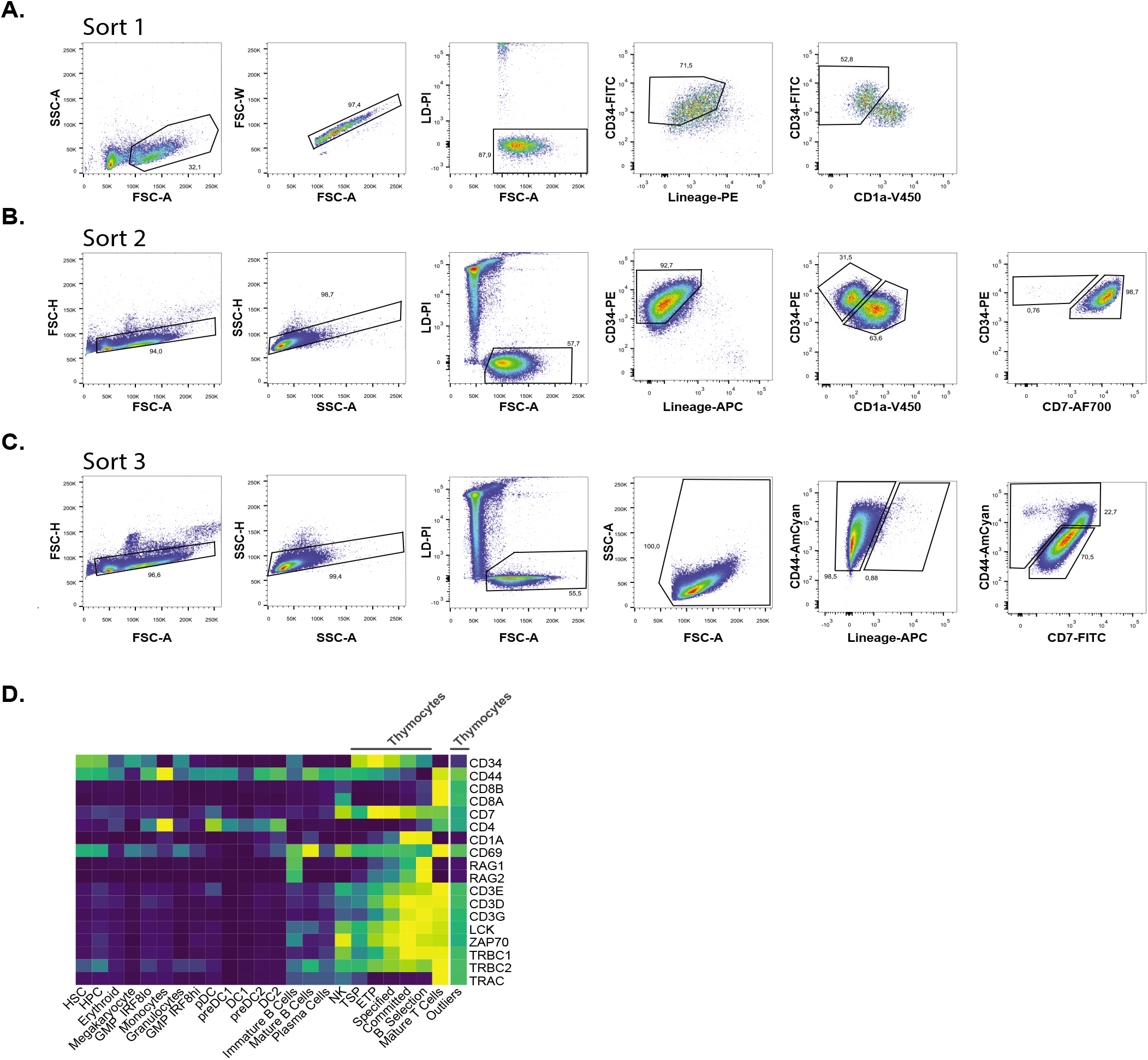
Validation of the sort and annotation quality. **A-C.** Gating strategy representative for each sort layout, used to isolate either viable lin^-^CD34^+^CD1a^-^ thymocytes (sort 1) (**A.**), viable lin^-^CD34^+^CD7^-/int^, lin^-^CD34^+^CD1a^-^ and lin^-^CD34^+^CD1a^+^ thymocytes (sort 2) (**B.**), and viable lin^+^, lin^-^CD44^+^CD7^-/+^ and lin^-^CD44^int^CD7^+^ thymocytes (sort 3) (**C.**). **D.** Heatmap visualization of pseudo bulk log2 counts, averaged and scaled gene-wise, identifying contaminating thymic mature T cells (outliers), prior to removal, compared to other developmental subsets present in either the BM, thymus or both datasets.

**Figure S2:**
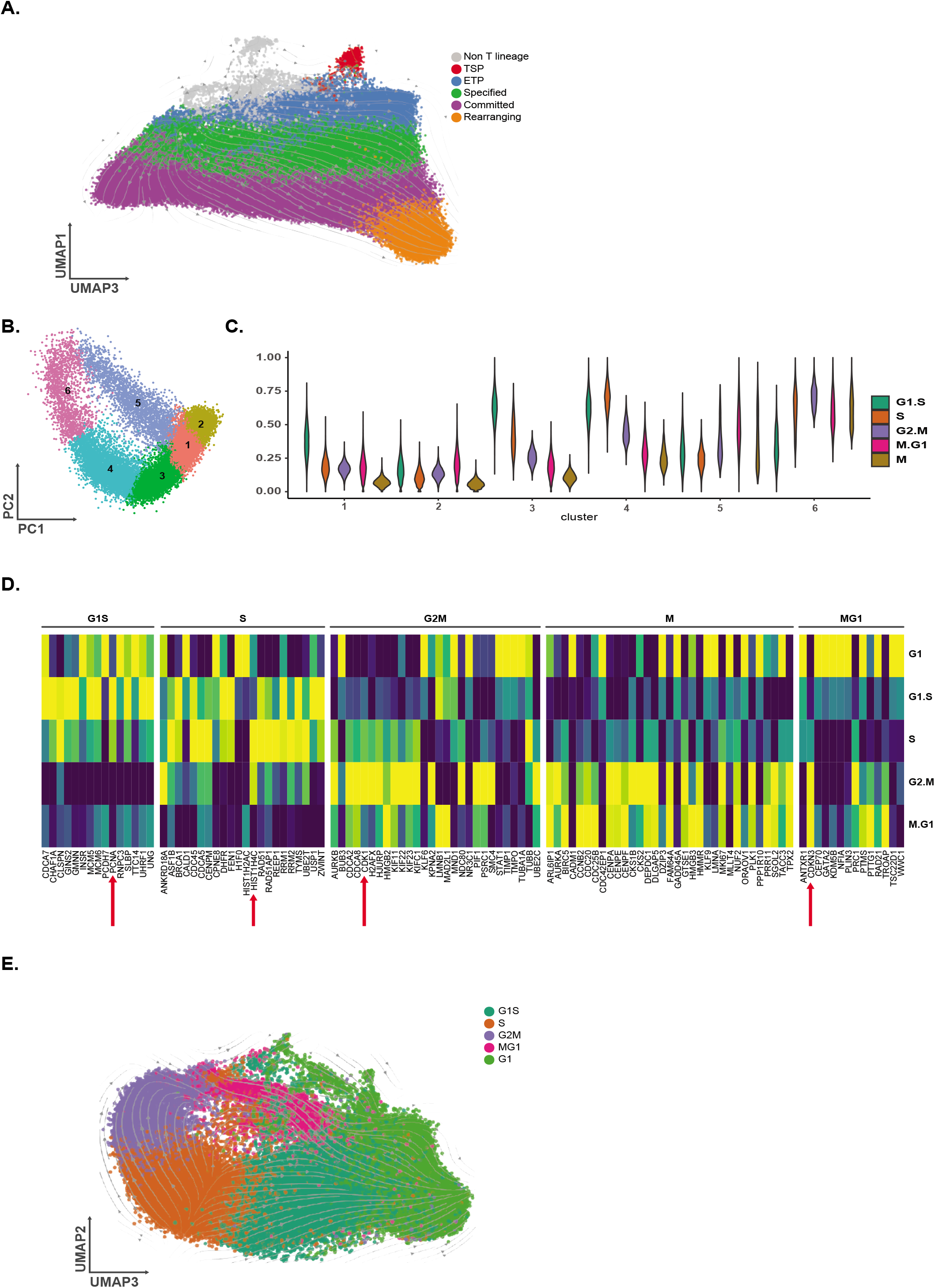
Development inferred by RNA velocyto and manual annotation of cell cycle stages on the lin-CD34+CD1a-fraction. **A.** Transition probabilities inferred by the RNA velocyto algorithm overlayed on the annotated populations displayed on dimension 1 & 3. **B.** MNN corrected PCA of lin^-^CD34^+^CD1a^-^ thymocytes using a curated set of cell cycle associated genes with the colors and numbers representing Leiden clustering. **C.** Violin plots depicting the averaged expression of all genes specific to a certain cell cycle stage (feature agglomeration), which were subsequently scaled. **D.** Heatmap of pseudo bulk log2 counts, averaged and scaled gene-wise, visualizing the expression of stage specific genes in each of the annotated cell cycle stages. Arrows represent the genes depicted in **Figure 2E**. **E.** Transition probabilities inferred by the RNA velocyto algorithm overlayed on the annotated cell cycle stages displayed on dimension 2 & 3.

**Figure S3:**
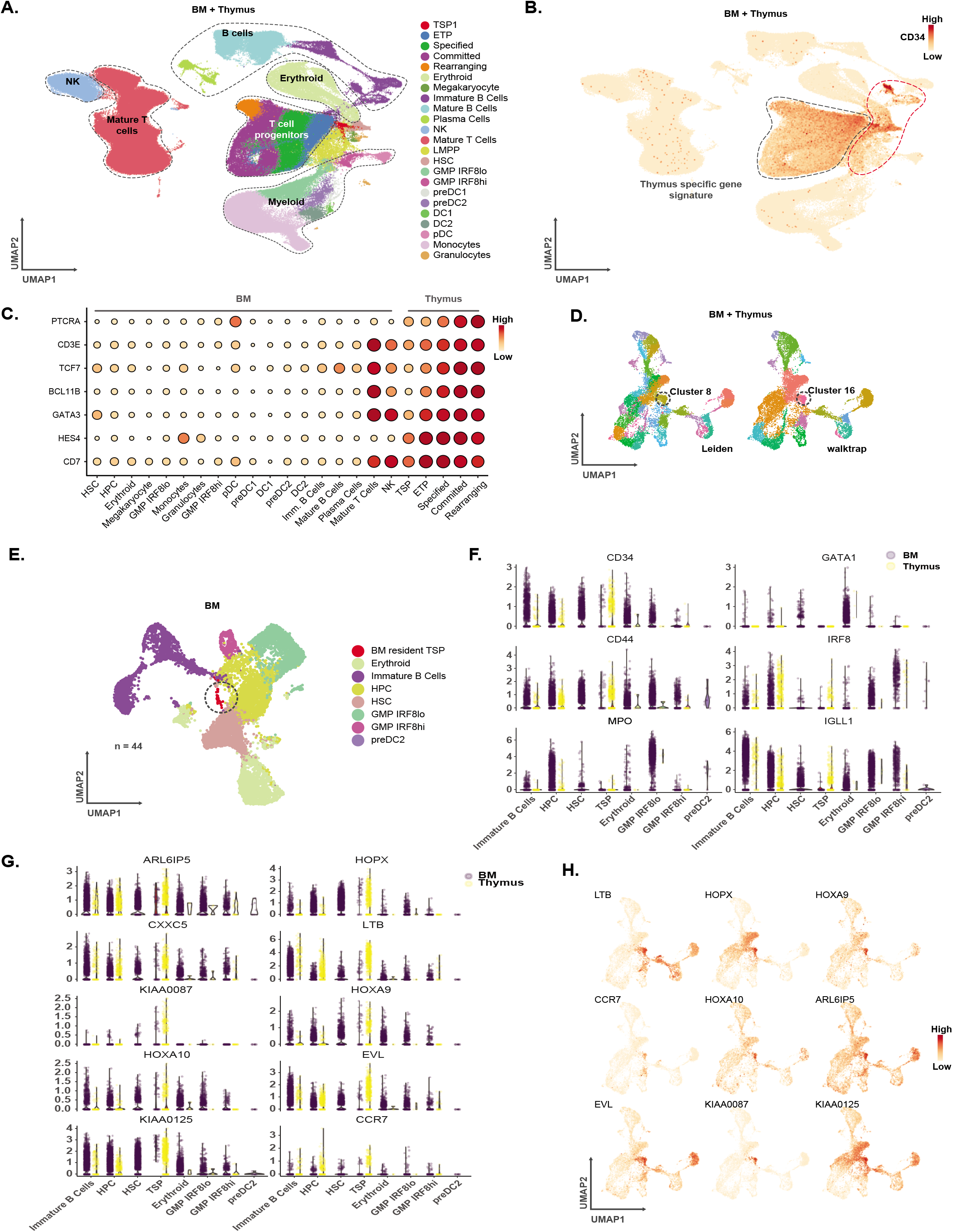
Identification and characterization of TSP across tissues. **A-C.** Isolation of CD34^+^ precursors following integration of the HCA BM and *ex vivo* thymocyte datasets. **A.** Annotated UMAP visualizing the subsets present in both datasets. **B.** Expression of CD34 in these subsets, with the isolated cells highlighted in grey, while excluding cells displaying co-expression of genes dependent on Notch signaling. **C.** Dot plot of pseudo bulk normalized log2 counts, averaged and scaled gene-wise, visualizing co-expression of genes dependent on Notch signaling used to exclude cells highlighted in grey **B**. The size of dots corresponds to gene expression. **D.** UMAP representing the integrated CD34^+^ BM cells and G1 thymocytes which lack a thymus specific signature visualizing both Leiden (left) and walktrap (right) clustering used to identify BM-resident TSPs which consistently cluster together. **E.** Annotated UMAP of CD34^+^ BM resident progenitors. **F-G.** Violin plots depicting gene expression signatures for thymocyte and BM progenitors separately (**F-G**), for genes described in literature (**F.**) and genes significantly upregulated in TSPs following a log ratio test (FDR <= 0.05; log fold change >= 0.5) (**G.**). **H.** UMAP of the integrated CD34+ BM and G1 thymocytes as described in **D.,** displaying significantly upregulated genes as described in **G**. The size of dots corresponds to gene expression levels.

**Figure S4:**
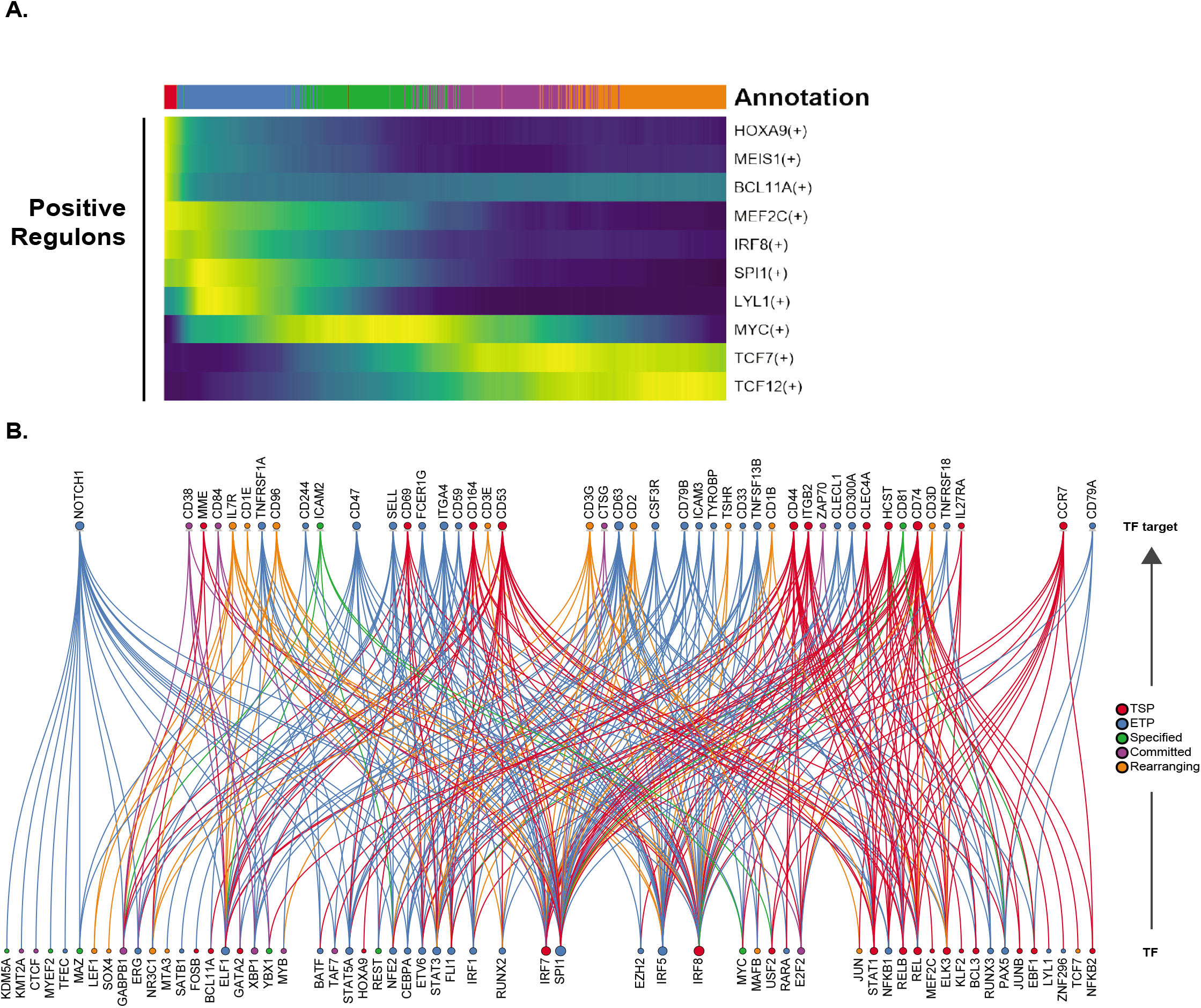
Validation of transcription factor activity and prediction of cell surface markers along pseudotime using SCENIC. **A.** Predicted TF activity as the area under the curve (AUC) of positive regulons associated with significantly overexpressed TF along pseudotime (FDR <= 0.05). **B.** Bipartite graph depicting significantly expressed TF during development (FDR <= 0.05) and their predicted binding on loci of TF. Node size reflects the number of connections between TFs and color is based on maximum of a loess curve fitted along pseudotime, through each gene.

**Figure S5:**
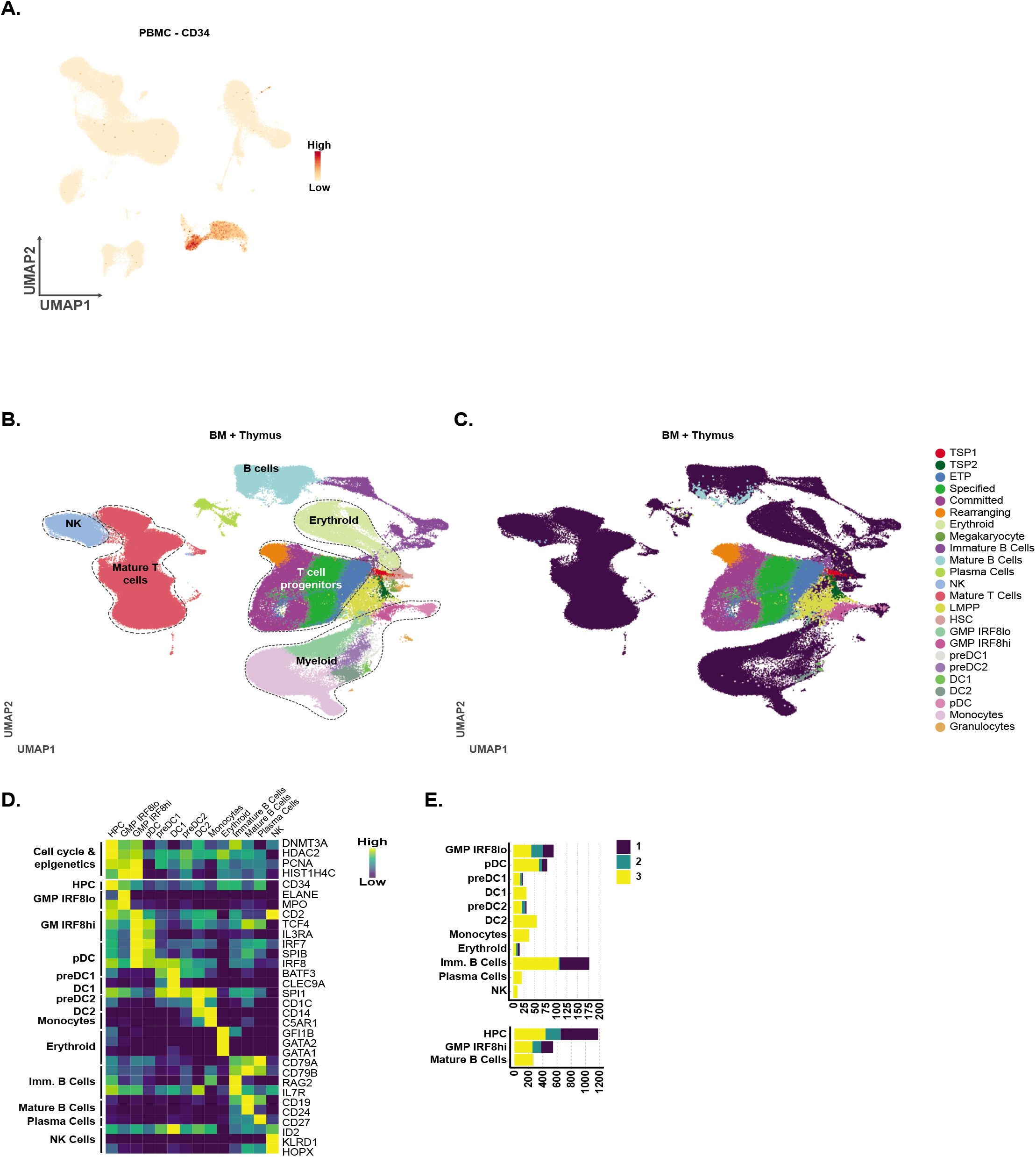
Identification and isolation of immature PBMCs and transcriptional heterogeneity of the non T-lineage CD34+ thymocytes. **A.** UMAP of human PBMCs visualizing CD34 expression, with an isolated fraction of immature cells highlighted in grey. The size of dots corresponds to gene expression levels. **B-C.** UMAP visualization of the integrated HCA BM and ex vivo human thymocyte datasets (sort 1-3) (**B-C.**), with all subsets annotated (**A.**) or only the subsets detected in the ex vivo human thymocyte dataset annotated (**B.**) (n > 3). **D.** Heatmap of the subsets detected in the ex vivo thymocyte dataset (sort 1-3) (n > 3) visualizing genes unique for each subset. **E.** Stacked bar charts displaying the number of detected cells per subset for each sorting strategy.

**Figure S6:**
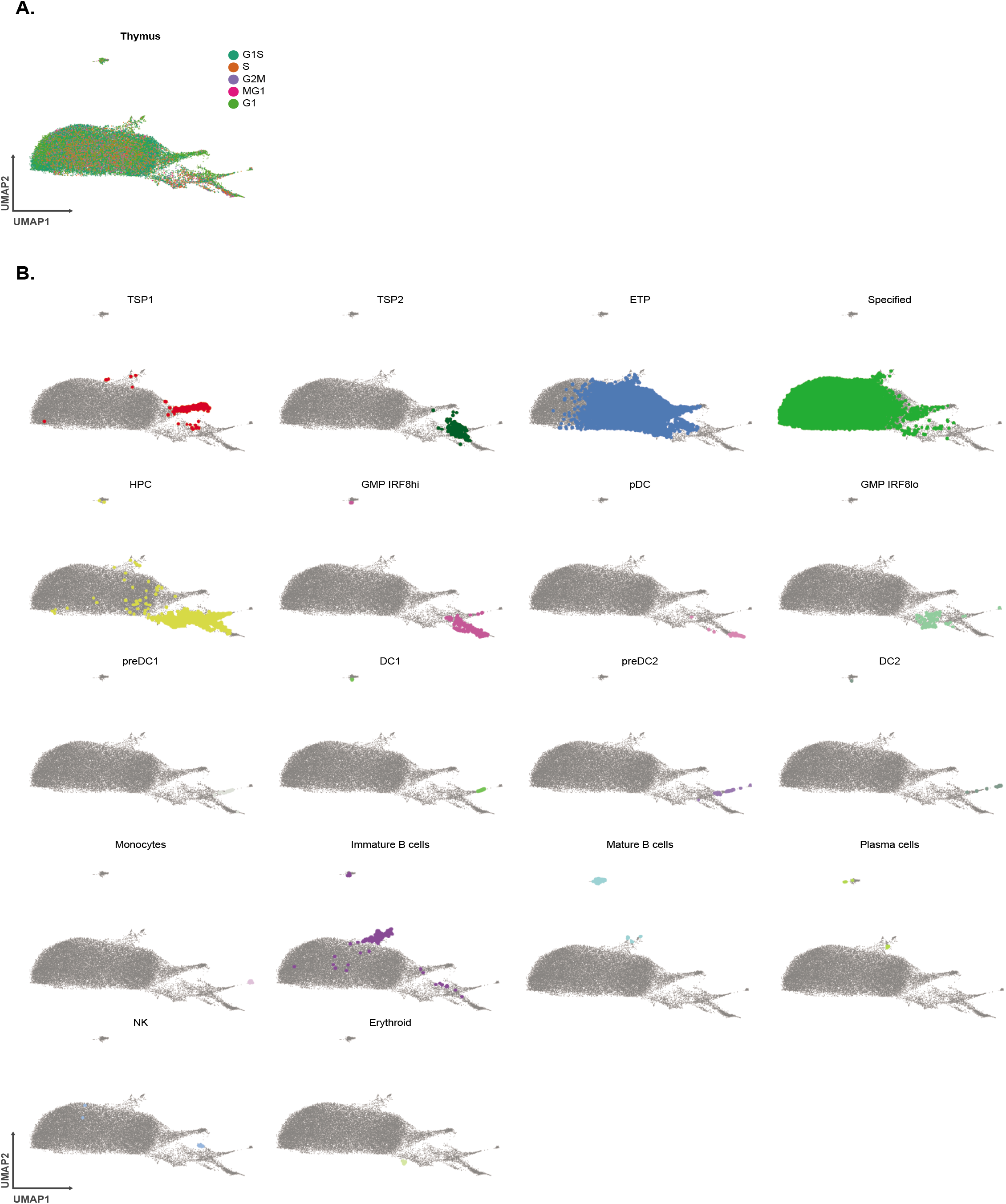
The heterogeneity in uncommitted CD34+ thymocytes is topologically reflected following cell cycle regression. **A-B.** UMAP visualization of uncommitted thymocytes following cell cycle regression, with assessment of the cell cycle stages (**A.**) and location of developmental subsets in the manifold (**B.**).

**Figure S7:**
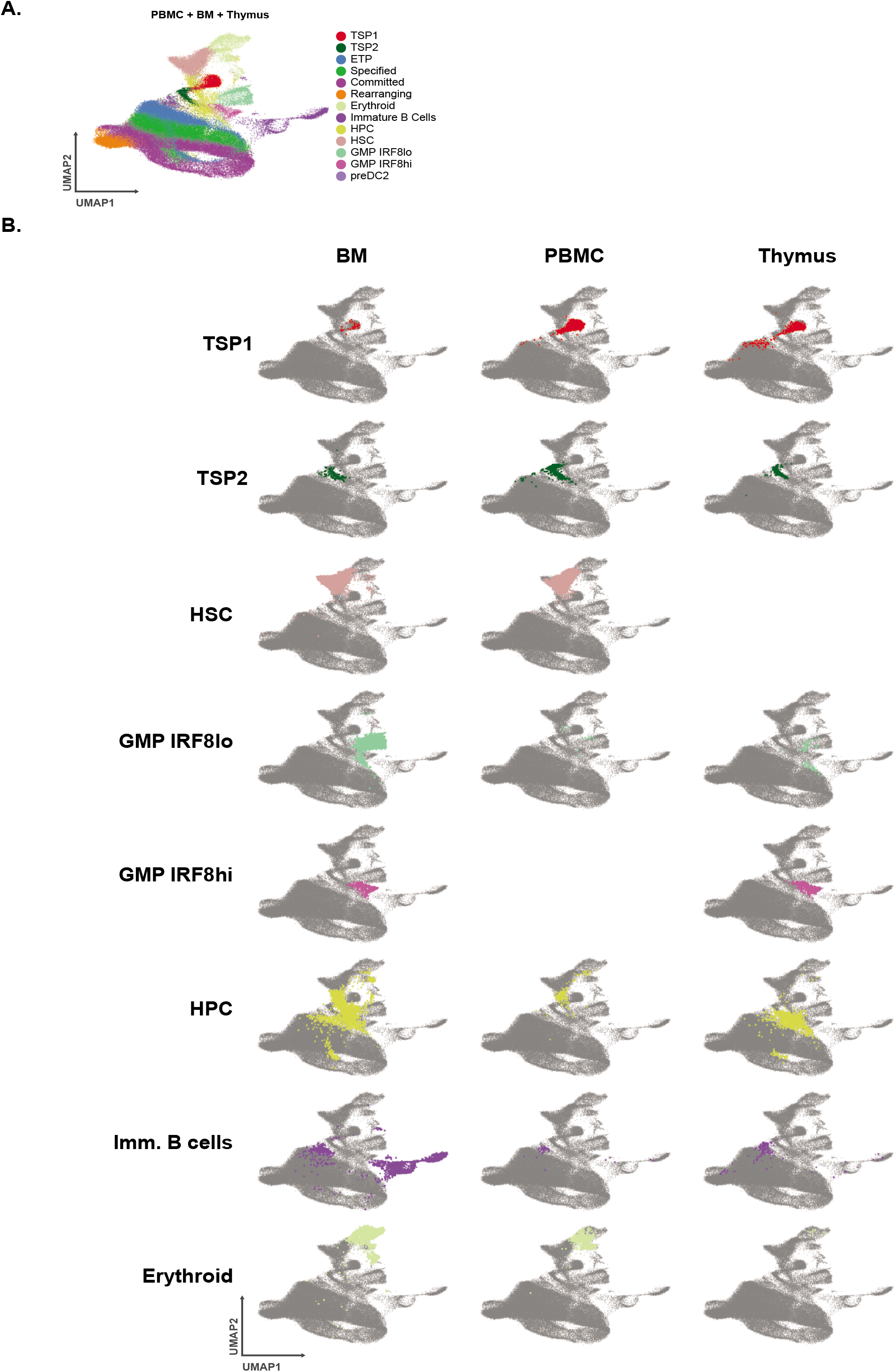
Validation of non T-lineage intrathymic development of plasmacytoid dendritic cells. **A-B.** UMAP visualizations of the integrated CD34^+^ BM progenitors, immature PBMCs and *ex vivo* thymocytes, excluding mature populations, with the included subsets annotated (**A.**) and visualized separately according tissue (**B.**).

